# CBL-C E3 ubiquitin ligase acts as a host defense to mediate ubiquitin-proteasomal degradation of enteropathogenic *Escherichia coli* Tir protein

**DOI:** 10.1101/2021.03.08.434331

**Authors:** Jinhyeob Ryu, Ryota Otsubo, Hiroshi Ashida, Tamako Iida, Akio Abe, Chihiro Sasakawa, Hitomi Mimuro

## Abstract

Translocated intimin receptor (Tir) is an essential bacterial factor for enteropathogenic *Escherichia coli* (EPEC) to establish Tir-intimin interaction-mediated adherence to the epithelial cell and to form actin pedestal structures beneath the adherent bacteria. However, it remains unclear how the host cells eliminate Tir protein after infection. Here we show that intracellular translocated Tir is degraded via the host ubiquitin-proteasome system. We found that host CBL-C, an E3 ubiquitin-protein ligase, bound to and ubiquitinated the 454 tyrosine-phosphorylated Tir protein. Tir ubiquitination leads to proteasome-dependent degradation and attenuated EPEC colonization of the epithelial cell. Using *Citrobacter rodentium*, a mouse model for EPEC, we demonstrated that infection with *C*. *rodentium* mutant expressing a tyrosine-phenylalanine-substituted Tir (CBL-C resistant) showed increased bacterial loads in the colon and lethality compared with infection with *C*. *rodentium* expressing wild-type Tir. These results indicate that CBL-C is a critical host defense factor that determines the fate of cytosolic Tir and terminates bacterial colonization.

**Graphical Abstracts:** 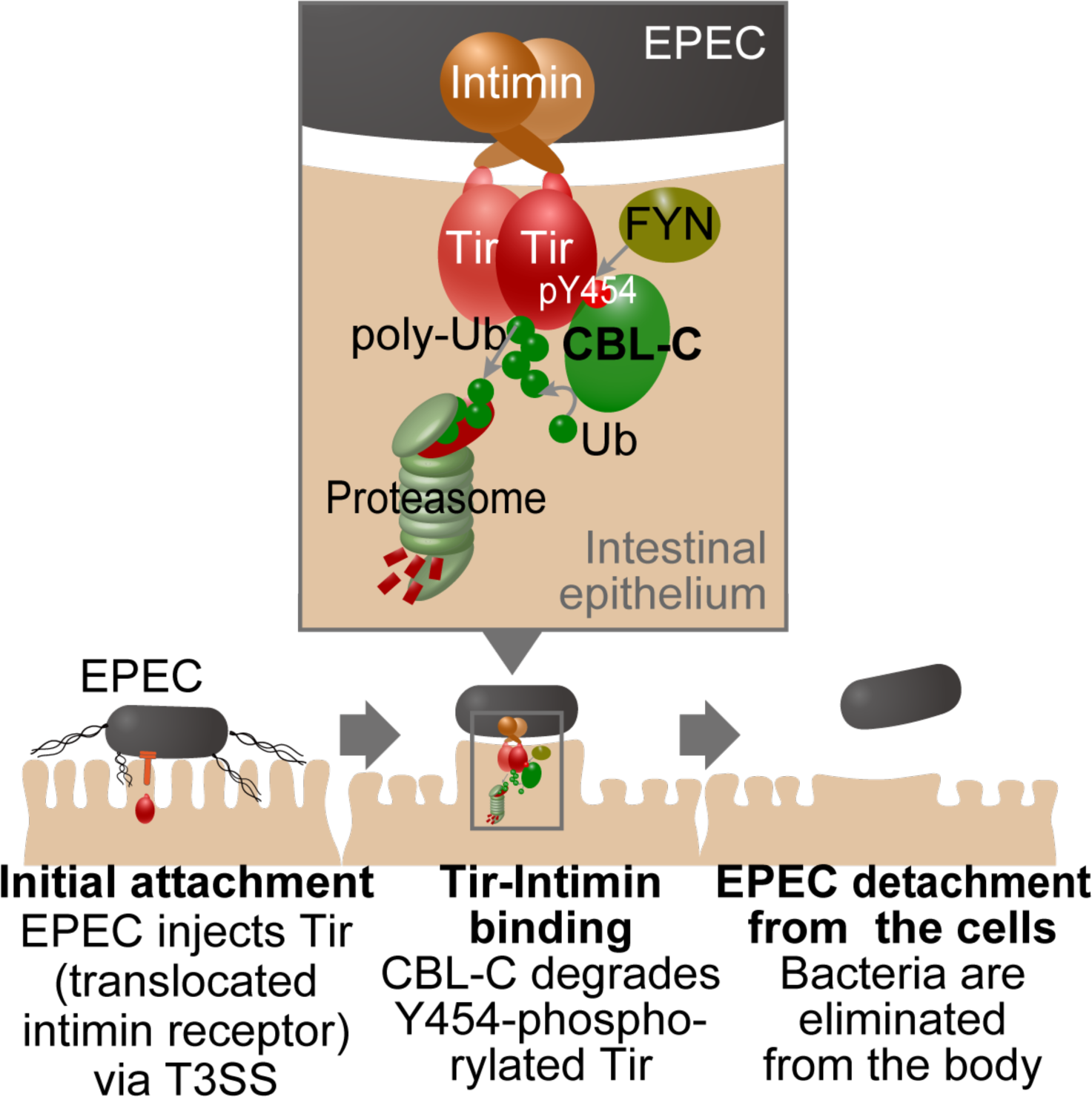

## Introduction

Enteropathogenic *Escherichia coli* (EPEC) is a human-specific pathogen that is the most prevalent diarrheal pathogen affecting infants and children, and causes acute and prolonged watery diarrhea. EPEC infection causes significant destruction of the tight junctions that constitute the intestinal epithelial barrier and massive imbalances of liquid and ions across the intestinal epithelial barriers, which leads to diarrhea.

EPEC is called an attaching-effacing (A/E) pathogen that effaces the microvilli and remodels host actin networks to form actin pedestal structures beneath the bacterium’s attachment site to the host cell (Stevens et al., 2006). Initial attachment of EPEC to the apical surface of an enterocyte is regulated by the bundle-forming pili (BFP). BFP extends from the bacterial cell surface to form unique “localized” adherence (Hicks et al., 1998). Subsequently, *<u>E</u>. coli*-<u>s</u>ecreted <u>p</u>roteins EspB and EspD are translocated into the host cell via the type III secretion system (T3SS), forming a pore structure, allowing the translocation of other effector proteins. Tir (translocated intimin <u>r</u>eceptor) protein, which functions as a receptor for the bacterial membrane protein Intimin (*eae*), is translocated to the bacterial attached cell via theT3SS (Deibel et al., 1998). Intimin binds and clusters Tir on the host cell surface, followed by EPEC delivery of effector proteins through the T3SS, and manipulation of host cellular signal transduction, which eventually leads to disruption of tight junctions, mitochondrial dysfunction, and alternations of golgi apparatus, microtubule formation, and inflammatory responses; which are the causes for severe inflammatory colitis (Wong et al., 2011).

When Tir binds to Intimin, the host tyrosine kinase FYN, a member of SRC family kinases (SFKs), induces tyrosine phosphorylation at the 474^th^ amino acid of Tir (Donnenberg et al., 1993; Kenny, 1999; Kenny and Warawa, 2001; Liu et al., 1999; Phillips et al., 2004; Swimm et al., 2004). Upon tyrosine phosphorylation, Tir recruits the host adaptor protein NCK, which binds neuronal Wiskott-Aldrich syndrome protein (N-WASP), activates the actin-related protein (Arp)2/3 complex, and induces actin polymerization and formation of the EPEC pedestal (Campellone et al., 2002; Vingadassalom et al., 2010; Wong et al., 2011). In addition, phosphorylation of Tyr 454 promotes NCK-independent, weak polymerization of actin, and promotes complex formation with phosphatidylinositol 3-kinase to produce phosphatidylinositol 3,4,5- triphosphate (Campellone and Leong, 2005; Sason et al., 2009). Other tyrosine-phosphorylation sites in Tir (Tyr483 and Tyr511) are reported to bind to SHP-1 phosphatase to inhibit TRAF6 ubiquitination which leads to suppression of immune response (Yan et al., 2012). Enterohemorrhagic *E. coli* (EHEC) and *Citrobacter rodentium* (*C. rodentium*) are also known to form pedestal-like structure, similar to EPEC. However, EHEC forms a pedestal-like structure via a signal independent of tyrosine phosphorylation of Tir. EHEC encodes an additional type 3-translocated effector, EspFU (also known as TccP) that binds and activates the host protein N-WASP to induce actin polymerization (Brady et al., 2007). Meanwhile, *C. rodentium* and EPEC Tir proteins are functionally compatible and both proteins require tyrosine phosphorylation to induce actin rearrangements for pedestal-like structure formation. Although the molecular mechanisms of pedestal-like structure formation during EPEC infection of epithelial cells are being well characterized, the fate of Tir protein transferred to the attached host cell remains unclear.

Ubiquitin (Ub) proteasome system (UPS) is an ATP-dependent proteolytic system for intracellular protein turnover and degradation. The protein substrates are marked for proteasome degradation by being linked to polypeptide cofactor Ub (a 76 amino acid peptide), through the cascade reactions of a series of enzymes: E1 (Ub-activating enzyme), E2 (Ub-conjugating enzyme), and E3 (Ub ligase enzyme). Ubiquitination has several binding patterns, which differ in their function (Walczak et al., 2012). Ub has seven lysine residues (Lys, K), and when forming polyubiquitin, the Lys and the Gly-76 at the C-terminal of another Ub form an amide bond. The function of Ub differs depending on which Lys is used for binding. In addition, N-terminal Met and C-terminal Gly may combine to form polyubiquitin. Polyubiquitin mediated by K48 transduces proteasome degradation signals. K63 polyubiquitin interacts with proteins to perform signal transduction and DNA repair. Linear polyubiquitin mediated by Met1 is involved in NF-κB activation (Walczak et al., 2012). These distinct Ub chains have different effects on the function of protein substrates.

CBL-C is a member of CBL (Casitas B-lineage Lymphoma) family of RING (Really Interesting New Gene) finger ubiquitin ligases. CBL family proteins are known to negatively regulate activated protein tyrosine kinases (PTKs) by ubiquitination of the targets PTKs for degradation either by assisting their endocytic sorting or by boosting proteasomal degradation (Mohapatra et al., 2013). CBL family proteins are composed of three mammalian homologues (CBL, CBL-B, and CBL-C) that are classified into two specific products: one is the longer CBL and CBL-B, which are highly conserved in their primary sequences as well as their domains; and the other is CBL-C, which is a shorter product that lacks some key C-terminal domains and motifs. All three members of the CBL family proteins share a highly homologous N-terminal region that serves as the tyrosine kinase binding (TKB) domain composed of four helix bundles (4H), an EF hand, and a SH2 domain (Meng et al., 1999). The TKB domain follows a Cbl linker sequence and a RING finger domain: these three N-terminal domains are important as structural platforms for interacting with E2 ubiquitin conjugating enzyme and are key to the E3 ubiquitin ligase activity of CBL family proteins (Lill et al., 2000; Ota et al., 2000; Zheng et al., 2000).

A previous proteomic screening study with 15-mer peptides showed that a peptide encompassing the phosphorylated Tyr454 region of Tir binds to SHP2, PI3K, and CBL-C. In contrast, a peptide containing the phosphorylated Tyr474 region of Tir binds to NCK1, NCK2, RasGAP1, and SRC (Selbach et al., 2009). Since CBL-C is an E3 Ub ligase, we investigated the relationship between the fate of Tir protein and CBL- C Ub ligase activity during EPEC infection of intestinal epithelial cells. We found that CBL-C binds to and degrades Tir protein in a UPS-dependent manner. Consistently, mice infection experiments with *Citrobacter rodentium* expressing Tir 474F (CBL-C–resistant Tir mutant) showed increased bacterial colonization in the mice intestine and more lethalithy than infection with *C. rodentium* expressing WT Tir (CBL-C–susceptible Tir). These results strongly suggest that CBL-C acts as a host defense system to terminate Tir activity and bacterial colonization of the intestinal mucosa.

## Results

### Proteasome-dependent protein degradation controls levels of translocated Tir

To determine the levels of Tir protein in EPEC-infected host epithelial cells, HCT116 human colon carcinoma cells were infected with a wild-type (WT) EPEC2348/60 at a MOI of 150. After 3 hours of infection, cells were treated with gentamicin and kanamycin to kill extracellular bacteria and the remaining cells were cultivated for another 1 to 3 hours. We then examined protein levels of Tir, Map and EspB by Western blotting. The levels of Tir protein in infected cells showed a time-dependent decrease to 10% of the sample at 3 hours after addition of gentamicin and kanamycin compared to baseline. In contrast, no time-dependent change was seen in the levels of Mitochondrial-associated protein (Map) or EPEC-secreted protein B (EspB) (Figure 1A).

**Figure 1.**
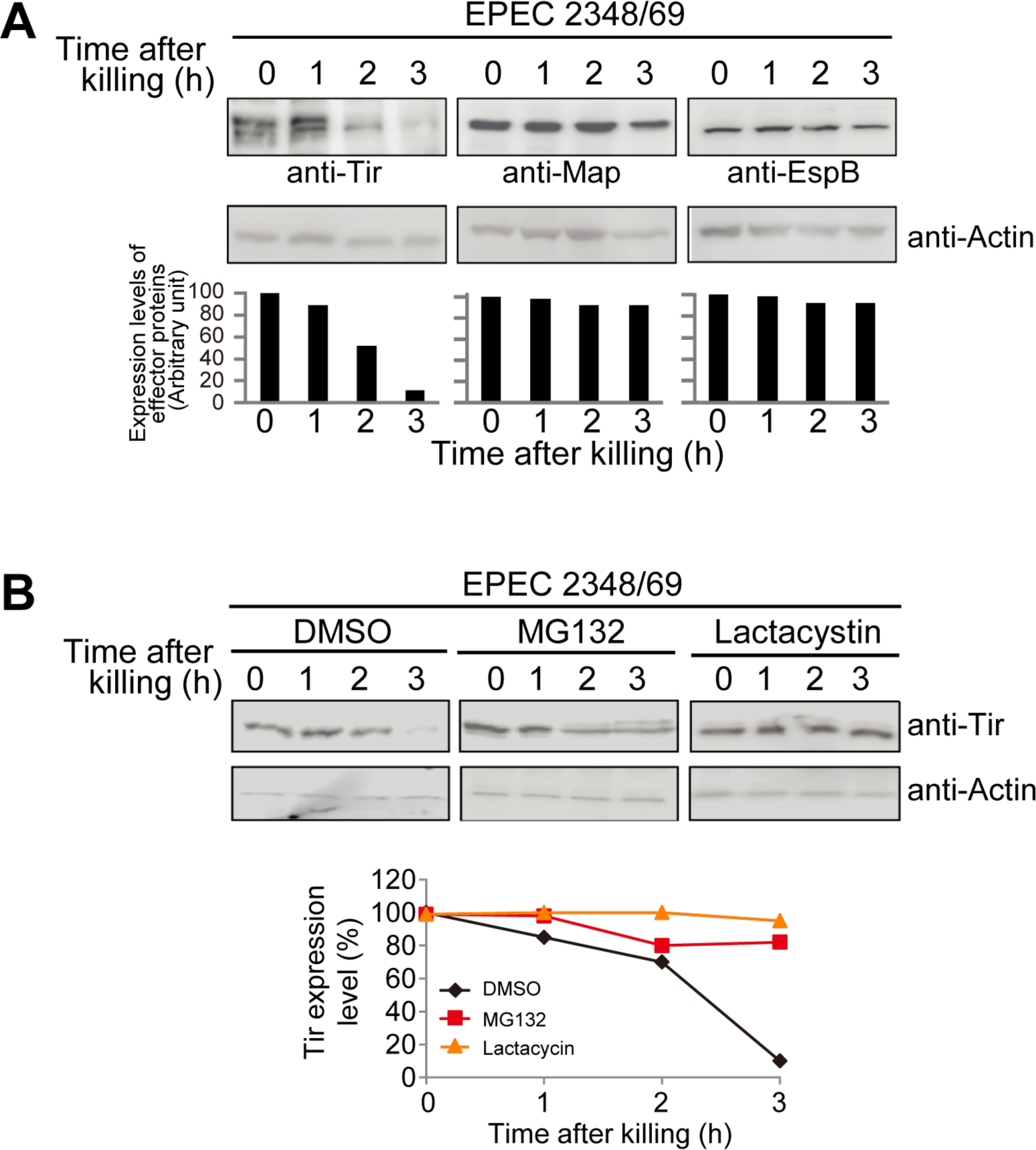
Proteasome-dependent protein degradation controls the levels of translocated Tir. **(A)** EPEC-injected Tir exhibit low stability within the host cells. HCT116 cells were infected with EPEC2348/69 for 3 hours. After washing, the cells were transferred into medium containing gentamicin and kanamycin to kill extracellular bacteria. After incubation for the indicated time, cells were harvested and lysates were separated by SDS-PAGE and subjected to western blot with the noted antibodies. **(B)** Proteasome inhibitors inhibit intracellular Tir protein degradation. The infected cells were treated with DMSO, MG132 (20μM) or Lactacystin (20μM) for the indicated time. Cells were harvested and lysates separated by SDS-PAGE and subjected to western blot with the noted antibodies. The intensity of proteins was normalized to the intensity of β-actin (Actin) and calculated in arbitrary units set to a value of 100 % for 0 h after killing. These results represent typical examples of three separate experiments. Densitometric analysis was conducted using ImageJ software.

There are two representative cellular protein degradation mechanisms in eukaryotic cells, Ubiquitin-Proteasome System (UPS), which degrades the majority of proteins, and autophagy, primarily responsible for the degradation of large aggregated proteins and damaged cellular organelles (Lilienbaum, 2013). To test if the degradation of Tir during infection is UPS-dependent, HCT116 cells were treated with MG132 or Lactacystin (proteasome inhibitors) and were infected with an EPEC strain. As shown in Figure 1B, the addition of the proteasome inhibitors rescued Tir from degradation, and the original levels remained constant for up to 3 hours after the addition of kanamycin and gentamicin to kill cell-attached EPEC, suggesting that Tir underwent UPS-dependent degradation.

### CBL-C targets Tir in a tyrosine phosphorylation-dependent manner

After attachment to the cellular surface, EPEC injects Tir protein into the host cell, and translocated Tir is phosphorylated by FYN tyrosine kinase (Phillips et al., 2004). As 15-mer Tir peptides containing 454 tyrosine-phosphorylated NPYAE- sequence bound to PI3K, SHP2, and CBL-C (Selbach et al., 2009), we tested if Tir has the capacity to bind CBL-C. To this end, we performed Glutathione S-transferase (GST) pull-down assays to identify the interaction between CBL-C and Tir with or without FYN kinase. As shown in Figure 2A, GST-CBL-C beads pulled down Myc-tagged Tir when FYN kinase was co-expressed in the cells, in which the Tir proteins were tyrosine phosphorylated. Similar results were obtained using GST-Tir beads (Figure 2B). As CBL-C is composed of Tyrosine Kinase Binding (TKB) region, of which Gly 276 to Glu (G276E) mutant of CBL-C (CBL-C GE) lost binding ability with PDGFR and EGFR, and a RING Finger (RF) region, of which Cys 351 to Ala (C351A) mutant (CBL-C CA) lost E3 ligase activity, and C-terminal region (Kim et al., 2004; Thien and Langdon, 2001), we created CBL-C GE, CBL-C CA, and CBL-C ΔC, which lack RF and C-terminal regions, and investigated which of the regions were responsible for the binding of CBL-C to Tir (Figure 2C). FLAG-tagged CBL-C WT, CA, and ΔC but not the GE mutant was co-immunoprecipitated with GST-tagged Tir from lysates expressing FYN (Figure 2D). Similar results were obtained by GST pull-down assay (Figure 2E). These results strongly suggested that CBL-C can interact with tyrosine-phosphorylated Tir, where the Gly 276 in the TKB region of CBL-C was involved in the interaction with Tir.

**Figure 2.**
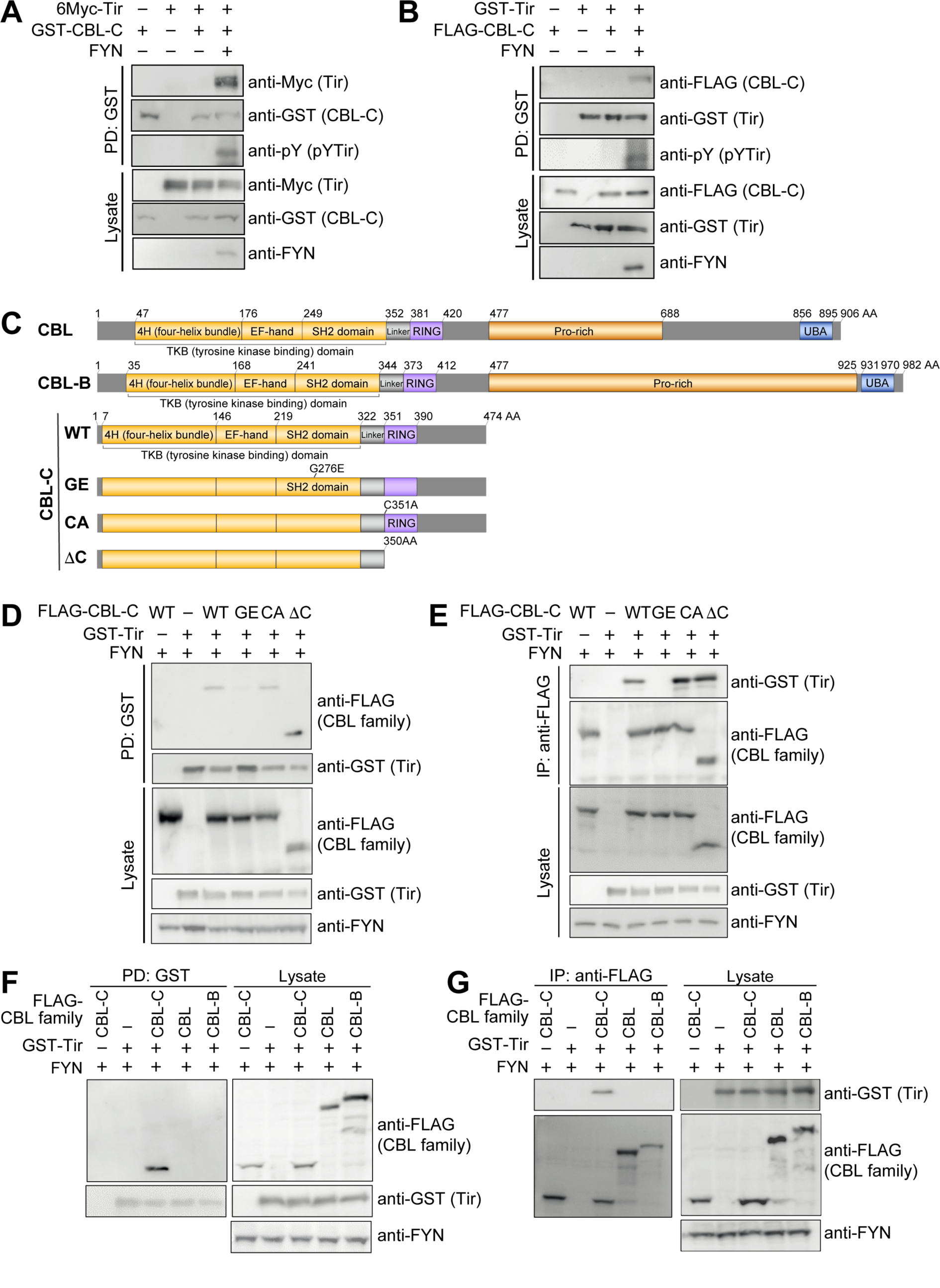
Tir binds to CBL-C in a tyrosine phosphorylation-dependent manner. **(A and B)** CBL-C binds to Tir in a phosphorylation-dependent manner. HEK293T cells were transiently transfected with the indicated plasmids. Proteins in the lysates were pulled down with Glutathione Sepharose 4B beads. Bound proteins were separated by SDS-PAGE and subjected to western blot with the noted antibodies. These results represent typical examples of three separate experiments. **(C)** Schematic diagram of CBL family proteins and CBL-C derivatives. **(D and E)** The Gly 276 in Tyrosine Kinase Binding domain of CBL-C is important for binding to Tir. HEK293T cells were transiently transfected with indicated plasmids. Proteins in the lysates were pulled down with Glutathione Sepharose 4B beads (D) or immunoprecipitated with anti-FLAG M2 and Protein G beads (E). Bound proteins were separated by SDS-PAGE and subjected to western blot with the noted antibodies. These results represent typical examples of three separate experiments. **(F and G)** Tir binds to CBL-C among CBL family. HEK293T cells were transiently transfected with indicated plasmids. Proteins in the lysates were pulled down with Glutathione Sepharose 4B beads (F) or immunoprecipitated with anti-FLAG M2 and protein G beads (G). Bound proteins were separated by SDS-PAGE and subjected to western blot with the noted antibodies. The results represent typical examples of more than three separate experiments. See also Figure S1.

The CBL family proteins (CBL, CBL-B, and CBL-C) are known to be products of proto-oncogenes (Swaminathan and Tsygankov, 2006). Knockout mice experiments indicated that while CBL and CBL-B play key physiological roles in tumor suppressors and autoimmune disease, the biological function of CBL-C remains unknown (Chiang et al., 2000; Griffiths et al., 2003; Murphy et al., 1998). Expression patterns indicated that CBL-C is highly expressed in intestinal epithelial cells, while CBL and CBL-B are expressed ubiquitously (Thien and Langdon, 2001). Amino acid sequence comparison revealed 82% identity in the TKB region between CBL and CBL-B, while 47% between CBL-C and CBL suggests that CBL-C has different characteristics compared to CBL or CBL-B (Figure S1). To investigate this selective binding capacity within the CBL family, we performed an immunoprecipitation assay as well as a GST-pull down assay. As shown in Figure 2F, FLAG-tagged CBL-C, but not CBL or CBL-B was co-immunoprecipitated with GST-tagged Tir from lysates expressing FYN. Similar results were obtained by GST pull-down assay (Figure 2G). These results indicated that host CBL-C protein specifically targets Tir among CBL family proteins.

### CBL-C induces Tir ubiquitination and degradation by the Ubiquitin-Proteasome System

To elucidate whether the E3 ligase activity of CBL-C is directly involved in Tir degradation, we performed *in vitro* ubiquitination assays using cell lysates containing HA-tagged Tir, His-tagged Ub, FYN, and FLAG-tagged CBL family (CBL-C, CBL, or CBL-B); and FLAG-tagged CBL-C CA (defective in E3 ligase activity) as a negative control. As shown in Figure 3A, Tir was ubiquitinated upon the co-expression of CBL- C, but not CBL-C CA, CBL, or CBL-B. It has been reported that K63-linked ubiquitination is related to lysosomal protein degradation, while K48-linked ubiquitination leads to UPS-dependent degradation (Grice and Nathan, 2016). We confirmed that K48-linked, but not K63-linked, ubiquitinated substrates were incorporated in HA-tagged Tir expressed in HEK293T cells (Figure 3B). The presence of the proteasome inhibitor MG132 showed increased levels of ubiquitinated Tir (Figure 3C). A cycloheximide (CHX) chase assay using HEK293T cells transfected with HA-Tir, FLAG-CBL-C and FYN confirmed that the presence of the proteasome inhibitor MG132 or Lactacystin rescued Tir from immediate degradation (Figures 3D and S2). These results indicated that the decreased levels of cytosolic Tir in host cells are caused by UPS-dependent degradation.

**Figure 3.**
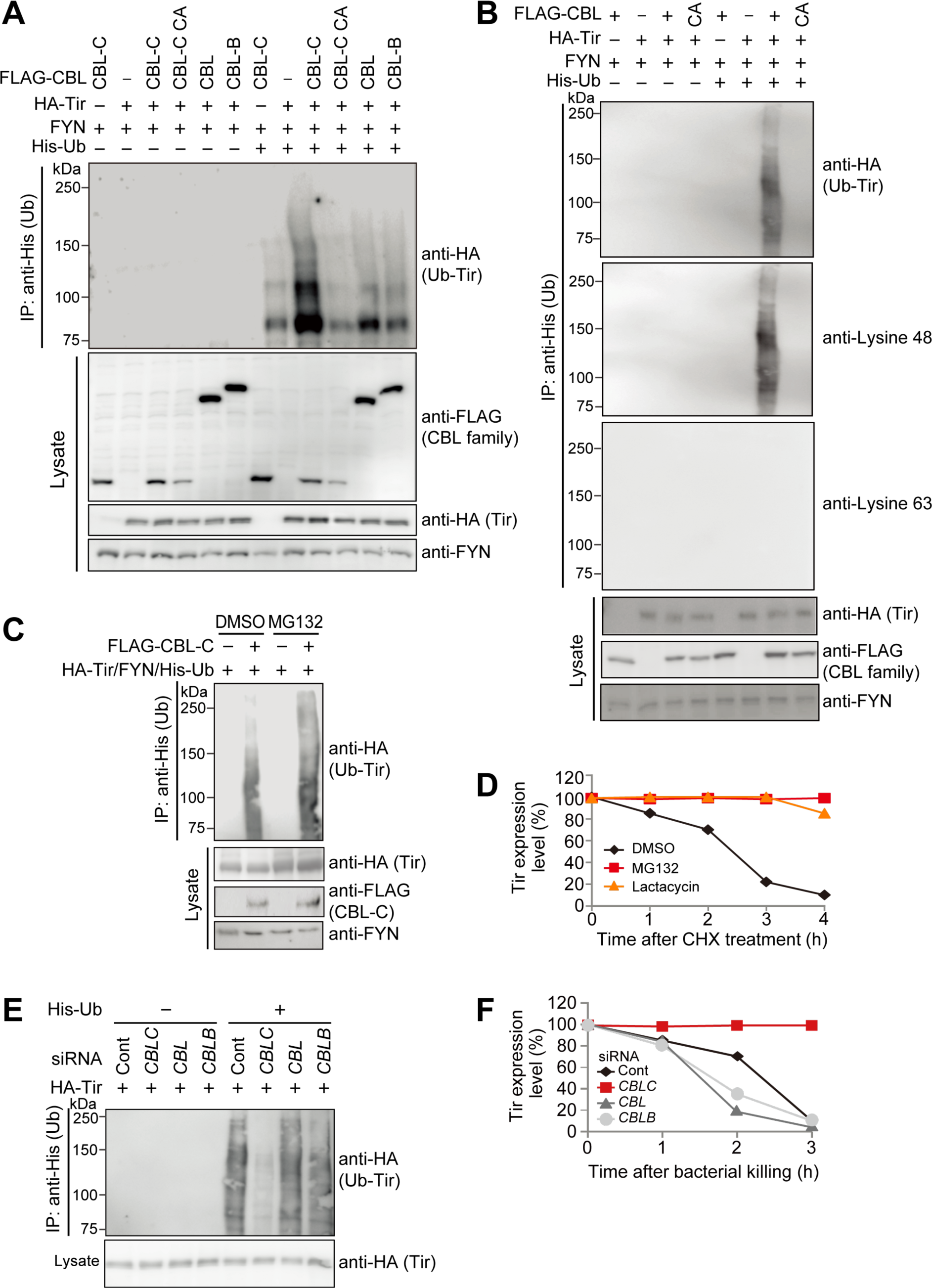
CBL-C induces Tir ubiquitination and degradation by the Ubiquitin-Proteasome System. **(A)** CBL-C induces Tir ubiquitination. HEK293T cells were transiently transfected with indicated plasmids. Proteins in the lysates were pulled down with Ni-NTA beads. Bound proteins were separated by SDS-PAGE and subjected to western blot with the noted antibodies. These results represent typical examples of three separate experiments. **(B)** CBL-C induces Tir Lysine 48 poly ubiquitination. HEK293T cells were transiently transfected with the indicated plasmids. Proteins in the lysates were pulled down with Ni-NTA beads. Bound proteins were separated by SDS-PAGE and subjected to western blot with the noted antibodies. **(C)** Proteasome inhibitor increases the accumulation of ubiquitinated Tir. HEK293T cells expressing the indicated plasmids were incubated with DMSO or MG132 (20 μM) for 4 hours. Proteins in the lysates were pulled down with Ni-NTA beads. The bound proteins were separated by SDS-PAGE and subjected to western blot with the noted antibodies. **(D)** UPS inhibitors attenuate Tir degradation. HEK293T cells were transfected with indicated plasmids, and treated with DMSO, MG132 (20 μM) or Lactacystin (20 μM) together with 10 μM of cycloheximide (CHX). After incubation for the indicated time, the lysates were separated by SDS-PAGE and subjected to western blot with the noted antibodies. The intensity of Tir protein was normalized to the intensity of β-actin (Actin) and calculated in arbitrary units set to a value of 100 % for 0 hour. The results represent the typical examples of three separate experiments. Densitometric analysis was conducted using ImageJ software. **(E)** Tir ubiquitination is medicated by CBL-C. HCT116 cells were transfected with siRNA targeting CBL family or Luc (control), then transfected with the indicated plasmids. The lysates were pulled down with Ni-NTA beads. Bound proteins were separated by SDS-PAGE and subjected to western blot with the noted antibodies. **(F)** HCT116 cells transfected with indicated siRNA targeting *CBL* family or Luc (control) were infected with EPEC Δ*tir*/*tir* WT at a MOI of 150 for 3 hours. The cells were washed 3 times with PBS, and then transferred into fresh medium containing gentamicin and kanamycin to kill extracellular bacteria. After incubation for the indicated time, cells were harvested and lysates separated by SDS-PAGE and subjected to western blot with the noted antibodies. The intensity of Tir protein was normalized to the intensity of β-actin (Actin) and calculated in arbitrary units set to a value of 100 % for 0 hour after killing. See also Figure S2 and S3.

To clarify the distinctive role of CBL-C among CBL family proteins in Tir degradation, we performed siRNA-mediated knockdown of each *CBL* family gene (Figure S3A). Silencing of *CBLC*, but not those of *CBL* or *CBLB*, showed significantly decreased levels of ubiquitination of intracellular Tir in HCT116 cells (Figures 3E and S3B). HCT116 cells treated with siRNA combinations specific for *CBL* family (*CBLC*, *CBL*, and *CBLB*) showed that Tir ubiquitination was rescued in cells overexpressing *CBL-C* but not *CBL* or *CBL-B* (Figure S3C). The specificity of CBL-C as an E3 ligase for Tir was also confirmed by assessment of Tir degradation levels within cells that were treated with siRNA combinations specific for *CBLC* and *CBL*, *CBLC* and *CBLB*, *CBL* and *CBLB*, and *CBL* family (*CBLC*, *CBL*, and *CBLB*); and by rescue experiments (Figures 3F, S3D, and S3E). Overexpression of CBL family proteins respectively in HCT116 cells treated with siRNA combinations specific for each of the *CBL* family members (*CBLC*, *CBL*, and *CBLB*) showed that degradation of Tir was rescued in HCT116 cells overexpressing *CBL-C* but not *CBL* or *CBLB-B* (Figure S3F). These data corroborated that the levels of Tir in host cells are regulated by CBL-C-dependent ubiquitination and degradation.

### CBL-C targets Tir Tyr 454 to degrade intracellular Tir Protein

A prior study reported that peptides corresponding to phosphorylated Tir Tyr 454 bind CBL-C protein (Selbach et al., 2009). To determine whether phosphorylation of Tir Tyr 454 is required for the CBL-C interaction, we generated a series of Tir derivatives in which the major tyrosine residues (Tyr 454 and Tyr 474) were substituted with phenylalanine (Y454F and Y474F, respectively) (Figure 4A). Although FYN was previously reported to phosphorylate Tyr 474 of Tir (Phillips et al., 2004), our *in vitro* kinase assay indicated that Tyr 454 in Tir was the major phosphorylation site by FYN (Figure 4B). To identify whether or not phosphorylation is required for the interaction of Tir with CBL-C, we performed GST pull-down assays to identify the interaction between CBL-C and Tir derivatives. As shown in Figure 4C, GST-CBL-C beads did not pull down 6Myc-tagged Tir when Tir Y454F mutant was co-expressed in HEK293T cells. Similar results with HCT116 cell lysate were reproducibly obtained using GST-Tir beads (Figure S4A). To ensure Tir Y454 takes part in the Tir ubiquitination, we performed *in vitro* ubiquitination assay using HEK293T cell lysate containing HA-Tir, FLAG-CBL-C, FYN and His-Ub. Tir was ubiquitinated when co-expressed with Tir WT, but not Tir Y454F (Figure 4D). When we used HCT116 cell lysate containing HA-Tir and His-Ub, Tir was also ubiquitinated when co-expressed with Tir WT, but not Tir Y454F (Figure S4B). To confirm the decrease in levels of intracellular Tir were dependent on Tyr 454 phosphorylation, we performed CHX chase assay in HCT116 and HEK293T cells. As shown in Figure 4E and S4C, Tir degradation was delayed in HEK293T cells co-expressed with HA-Tir Y454F, FLAG-CBL-C, and FYN, but not Tir WT. When we used HCT116 cell lysate containing HA-Tir derivatives, Tir degradation was also delayed when co-expressed with Tir Y454F, but not Tir WT (Figure S4D). These data demonstrate that Tir residue Tyr 454 is the target of CBL-C.

**Figure 4.**
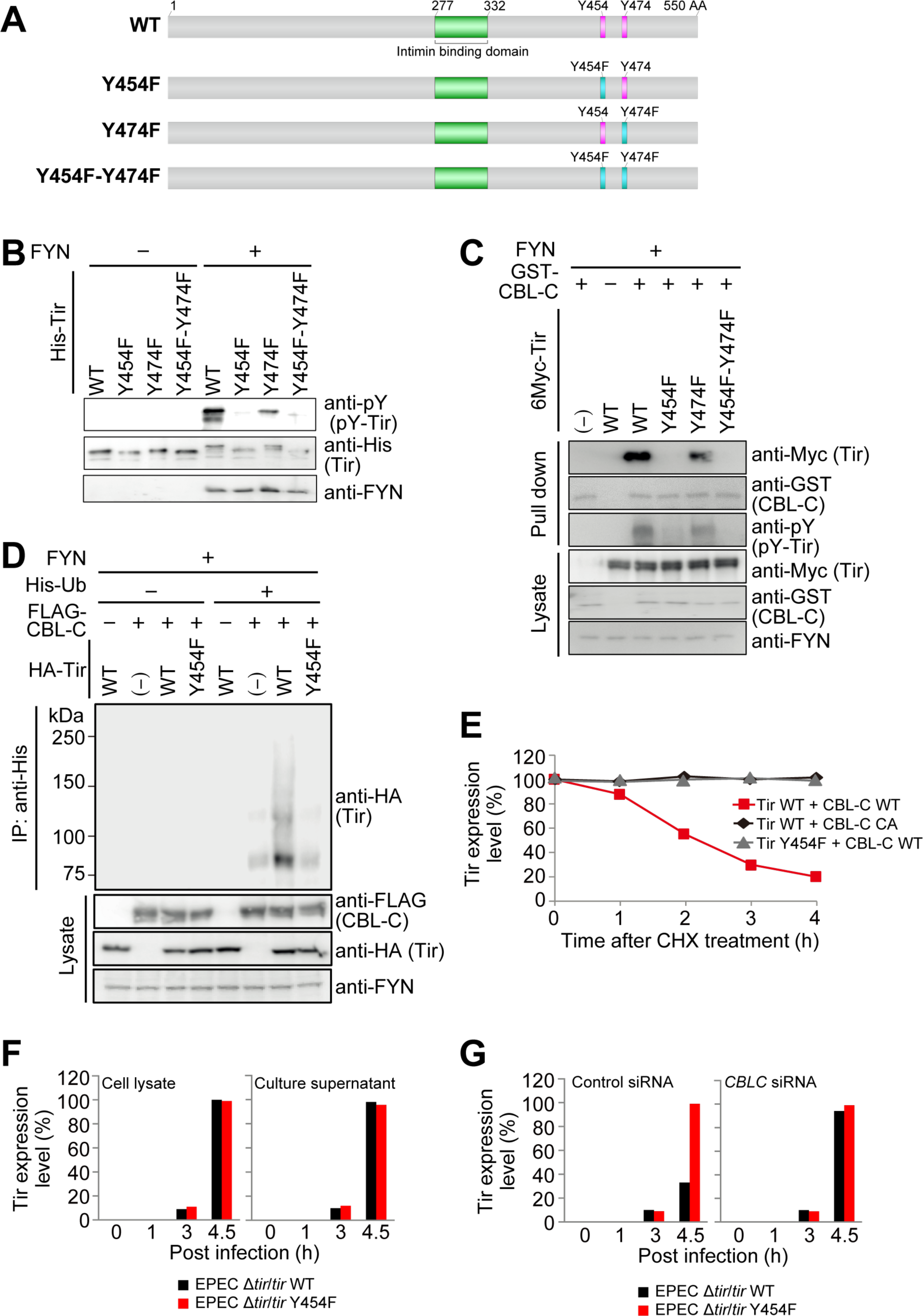
CBL-C targets Tir Tyr 454 to degrade intracellular Tir Protein. **(A)** Schematic diagrams of *Escherichia coli* O127:H6 (strain E2348/69 / EPEC) Translocated intimin receptor Tir. **(B)** Tir Tyrosine 454 is phosphorylated by FYN kinase. *In vitro* FYN kinase assays using His-Tyr WT or mutants as substrates. After incubation at 30°C for 20 min, the samples were separated by SDS-PAGE and subjected to Western blot with the noted antibodies. **(C)** CBL-C binds to phosphorylated Tir at residue Tyrosine 454. HEK293T cells were transiently transfected with indicated plasmids. The cells were washed 3 times with PBS and lysed for 30 minutes at 4°C in lysis buffer. The lysates were cleared by centrifugation and proteins were pulled down for 2 hours with Glutathione Sepharose 4B beads at 4°C. The bound proteins were washed 5 times with lysis buffer, separated by SDS-PAGE, and subjected to Western blot with the described antibodies. **(D)** Tir residue Tyrosine 454 is important for Tir ubiquitination. HEK293T cells were transiently transfected with the indicated plasmid combinations. Cells were washed 3 times with PBS and lysed in lysis buffer for 30 minutes at RT. Lysates were cleared by centrifugation, and proteins were pulled down with Ni-NTA beads for 2 hours at RT. Bound proteins were washed 5 times with lysis buffer, separated by SDS-PAGE, and subjected to Western blot with the indicated antibodies. These results represent typical examples of 3 separate experiments. **(E)** Tir residue Tyrosine 454 is important for Tir degradation. HEK239T cells expressing the indicated plasmids were treated with chcloheximide (CHX, 10 μM). After incubation for the indicated time, cells were harvested and lysates were separated by SDS-PAGE and subjected to Western blot with the indicated antibodies. The intensity of Tir protein was normalized to the intensity of β-actin (Actin) and calculated in arbitrary units set to a value of 100 % for 0 hour, These results represent typical examples of 3 separate experiments. Densitometric analysis was conducted using ImageJ software. **(F)** Y454F Tir protein levels in EPEC Δ*tir*/*tir* Y454F are equal to that in EPEC Δ*tir*/*tir* WT. Bacterial whole cell lysates (Bacteria) and culture supernatants (Medium) were separated by SDS-PAGE and subjected to western blot with the noted antibodies. The intensity of Tir protein was calculated in arbitrary units set to a value of 100 % for samples of EPEC Δ*tir*/*tir* WT at 4.5 hours. These results represent typical examples of three separate experiments. Densitometric analysis was conducted using ImageJ software. **(G)** CBL-C regulate protein levels of intracellular Tir secreted by EPEC. HCT116 cells transfected with indicated siRNA targeting *CBL* family or Luc (control) were subjected to additional transfection of CBL-C protein, and were infected with the indicated EPEC strains at a MOI of 150 for indicated time. Cell lysates were harvested and lysates were separated by SDS-PAGE and subjected to western blot with the noted antibodies. The intensity of Tir protein was normalized to the intensity of β-actin (Actin). The intensity of Tir protein was calculated in arbitrary units set to each value of 100 % for samples of EPEC Δ*tir*/*tir* Y454F at 4.5 hours. The results represent the typical examples of three separate experiments (mean and SEM). Densitometric analysis was conducted using ImageJ software. See also Figure S4.

To test the effect of Tir Tyr 454 phosphorylation on intracellular Tir protein levels, HCT116 cells were infected with EPEC Δ*tir* expressing wild type (WT) Tir (Δ*tir*/*tir* WT) or Y454F Tir (Δ*tir*/*tir* Y454F). Levels of Tir protein expression in bacteria as well as in secreted medium were comparable between EPEC Δ*tir*/*tir* WT and Δ*tir*/*tir* Y454F, at 1, 3, and 4.5 hours after incubation (Figures 4F and S4E). However, levels of intracellular-translocated Tir protein in control siRNA-treated HCT116 cells 4.5 hours after infection was decreased in EPEC Δ*tir*/*tir* WT infected cells, to a low level of about one third of that of EPEC Δ*tir*/*tir* Y454F infected cells (Figures 4G and S4F). This finding suggests that intracellular WT Tir but not Y454F Tir was degraded in host cells during EPEC infection. To demonstrate the effect of CBL-C on WT Tir degradation in infected cells, we prepared HCT116 cells in which CBL-C expression was blocked by siRNA. The results showed that the levels of intracellular Tir in cells treated with *CBLC* siRNA infected with EPEC Δ*tir*/*tir* WT were almost equal to that of EPEC Δ*tir*/*tir* Y454F (Figures 4G and S4F). As the levels of WT Tir were not rescued in EPEC Δ*tir*/*tir* WT-infected cells treated with control, *CBL* or *CBLB* siRNA (Figure S4G), these results further indicate that host cell CBL-C targets Tyr 454 residue of Tir to down-modulate the levels of intracellular Tir protein in cells infected with EPEC.

### Effect of Tir residue Tyr 454 and CBL-C on EPEC adherence to host cells

As it was previously reported that phosphorylate of Tir Tyr 474 is required for pedestal formation upon EPEC infection (Phillips et al., 2004), we evaluated the effects of Tyr 454 of Tir and its binding partner CBL-C on pedestal formation as well as bacterial attachment to host cells. HCT116 cells treated with control or *CBLC* siRNAs were infected with Δ*tir*/*tir* WT or Δ*tir*/*tir* Y454F of EPEC. After 4.5 hours of infection, bacterial attachment and actin pedestals (identified as actin clots beneath the adhered bacteria), were quantified by confocal microscopy (Figures S5A, S5B, 5A, and 5B). These results showed nearly two-thirds less bacterial adherence on control siRNA-treated cells by EPEC Δ*tir*/*tir* WT, as compared to those by EPEC Δ*tir*/*tir* Y454F. However, when cells were treated with *CBLC* siRNA, the number of adherent EPEC Δ*tir*/*tir* WT was almost comparable to that of adherent EPEC Δ*tir*/*tir* Y454F (Figures 5A and 5B), unlike when cells were treated with other *CBL* family siRNA (Figures S5C and S5D). The levels of pedestal formation showed the same tendency as bacterial adherence (Figures 5C and S5E). These data demonstrate that the interaction of Tir Tyr 454 and CBL-C affects the efficiency of EPEC adherence to host cells.

**Figure 5.**
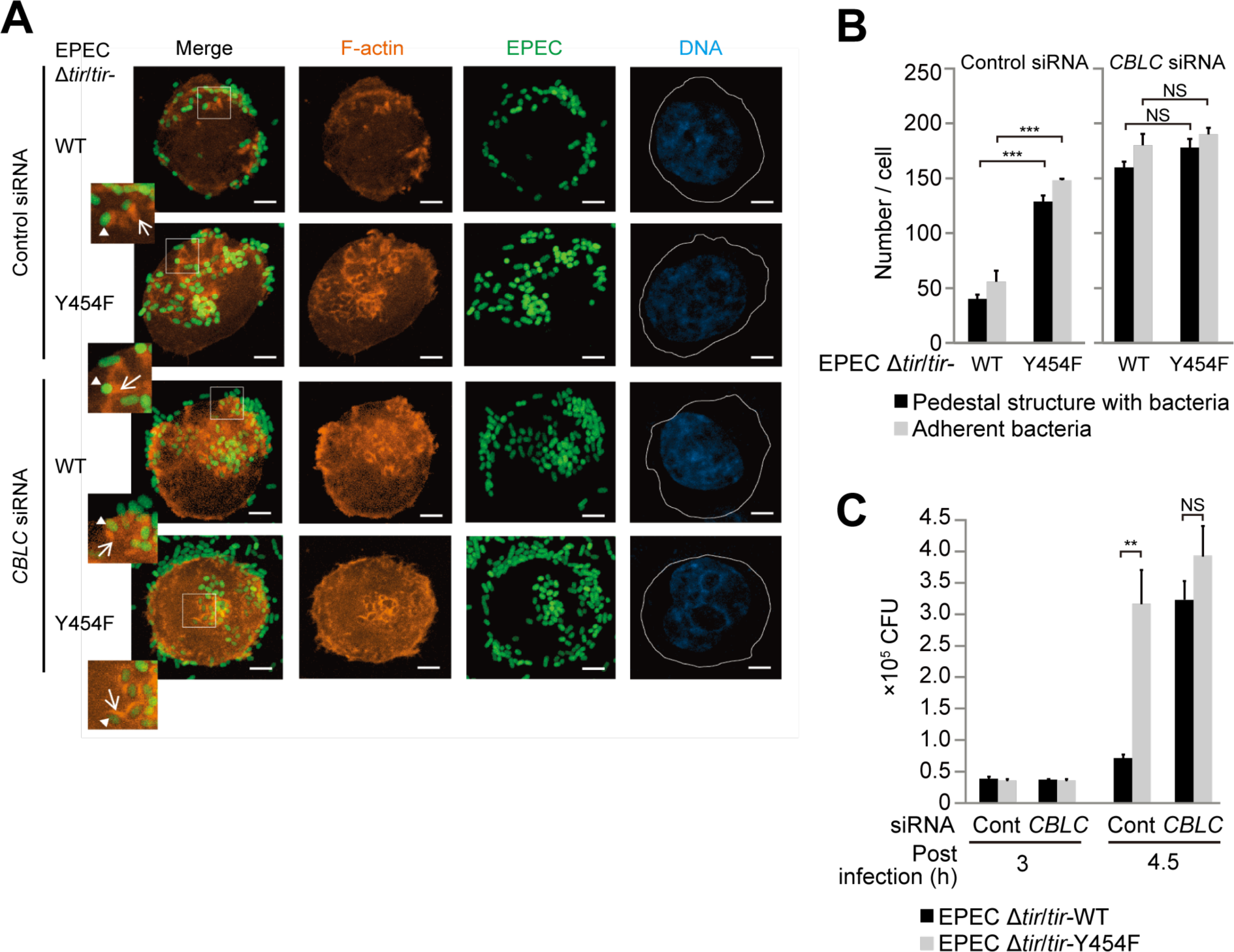
Effect of Tir residue tyrosine 454 and CBL-C on EPEC adherence to the host cells. (**A** and **B**) HCT116 cells transfected with siRNA targeting *CBL* family or Luc (control), or subjected to additional transfection of CBL-C protein were infected with the indicated GFP-EPEC strains at a MOI of 150 for 4.5 hours. The cells were fixed and actin (red), EPEC (green), and DNA (stained with DAPI, blue) were visualized. Scale bar, 5 µm (A). Quantification of pedestal structure and adhesion bacteria are shown (B). These results represent typical examples of three separate experiments. Data are represented as mean ± SEM (n = 150). N.S, not significant, \*\*\**P <* 0.001 (Student’s *t*- test). (**C**) HCT116 cells were transfected with siRNA of *CBLC* or Luc (control), and infected with the indicated EPEC strains at a MOI of 150 for the indicated time. The adhered bacteria were harvested at the indicated time points and plated on LB agar-plates to count CFUs. These results represent typical examples of three separate experiments. Data are represented as mean ± SEM (n = 3). \**P <* 0.01, \*\**P <* 0.005 and \*\*\**P <* 0.001 (Student’s *t*-test). See also Figure S5.

### CBL-C acts as a host defense

To establish the biological impact of CBL-C-mediated Tir degradation over EPEC infection, we employed the *Citrobacter rodentium* mouse infection model (Vallance et al., 2003). Since *C. rodentium* Tir (CrTir) shares amino acid sequence homology with EPEC Tir (Figure 6A), we tested whether CrTir Tyr 451 (equivalent to EPEC Tir Tyr 454) but not CrTir Y451F can undergoes UPS-dependent degradation by CBL-C. To this end, we performed an *in vitro* ubiquitination assay of HEK293T cell lysate containing FYN, FLAG-CBL-C, HA-Tir, and His-Ub. As shown in Figure 6B, CrTir WT, but not CrTir Y451F, was ubiquitinated in the presence of CBL-C. To ensure the decrease of intracellular CrTir resulted from UPS-dependent Tir degradation, we performed a CHX chase assay. As shown in Figure 6C, the HA-CrTir WT degradation was detected in lysate from HEK293T cells co-expressing FYN, FLAG-CBL-C, and HA-CrTir WT in the presence of CHX, which was canceled in lysate from HEK293T cells co-expressing FYN, FLAG-CBL-C, and HA-CrTir Y451F.

**Figure 6.**
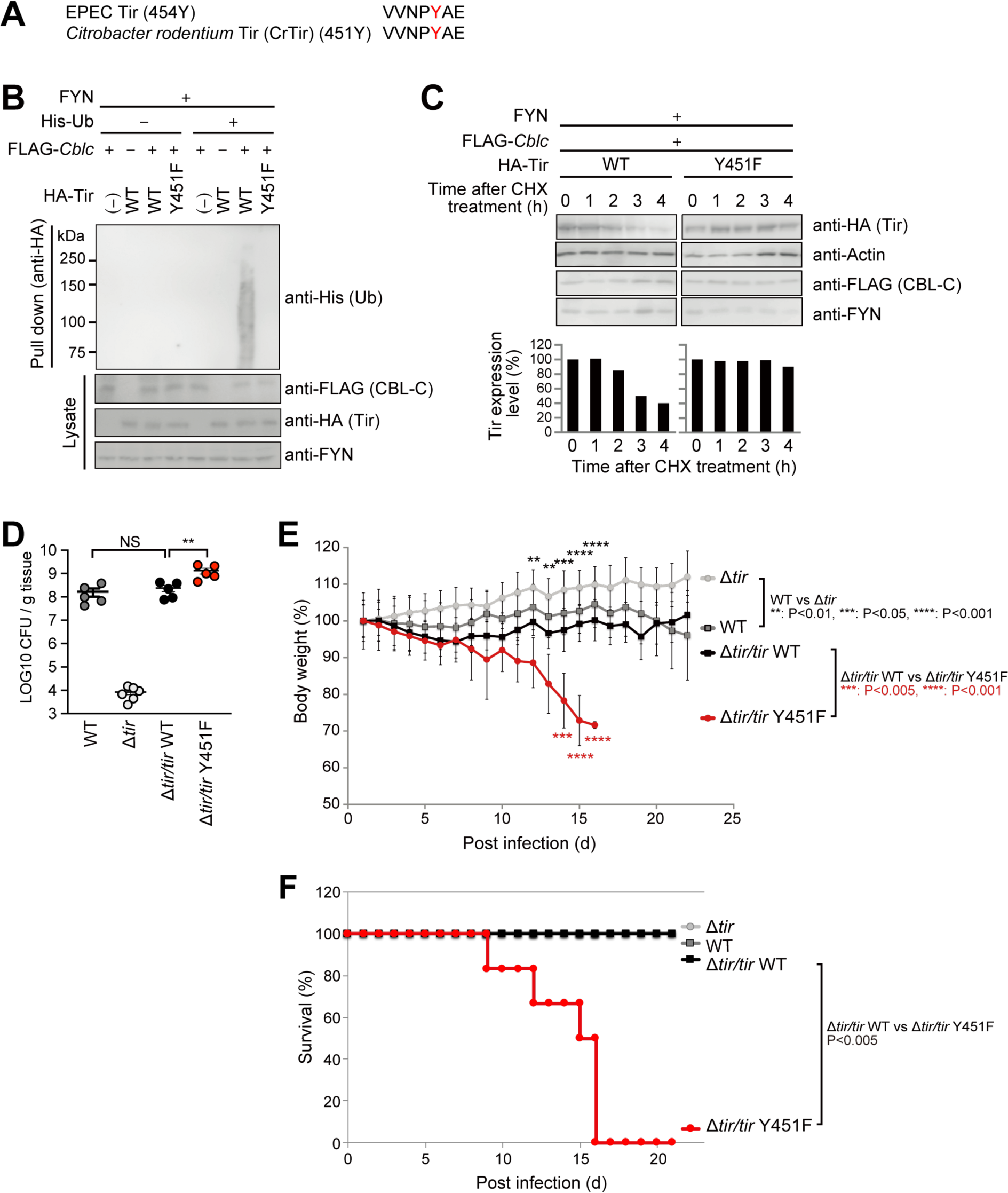
Pedestal structure and colonization are negatively regulated by CBL-C as a host defense. (A) Amino acid sequence alignment around EPEC Tir residue tyrosine 454 with *C. rodentium* Tir (CrTir) residue tyrosine 451. (B) HEK293T cells were transiently transfected with the indicated plasmids. The cells were washed 3 times with PBS and lysed for 30 min at RT in lysis buffer. The lysates were cleared by centrifugation, and proteins were pulled down for 2 hours with Ni-NTA beads at RT. Bound proteins were washed five times with lysis buffer, separated by SDS-PAGE and subjected to western blot with the noted antibodies. (C) HEK293T cells were transfected with the indicated plasmids, and treated with cycloheximide (10 μM). After incubation for the indicated time, cells were harvested and lysates were separated by SDS-PAGE and subjected to western blot with the noted antibodies. The intensity of Tir protein was normalized to the intensity of β-actin (Actin) and calculated in arbitrary units set to a value of 100 % for 0 hour. The results represent the typical examples of three separate experiments. Densitometric analysis was conducted using ImageJ software. (**D**–**F**) Studies of C3H/HeJ mice infected with *C. rodentium*. Six to 8-week-old C3H/HeJ mice were orally infected with *C. rodentium* strains (*C. rodentium* DBS100 strains WT, Δ*tir*, Δ*tir*/*tir* WT, and Δ*tir*/*tir* Y451F) and sacrificed 7-days post infection (*n* = 5 or 6 per group). CFU in colon was counted. Data are represented as mean ± SEM (D). C3H/HeJ mice were orally infected with *C. rodentium* strains and body weight was monitored daily up to 21 days. Percentage of mice body weight from 0 day after infection is shown. (*n* = 6 mice per group.) Data are represented as mean ± SEM (E). C3H/HeJ mice were orally infected with *C. rodentium* strains and survival rate was monitored daily up to 21 days. Percentage of mice survival from initial population is shown. (*n* = 6 mice per group). Statistical significance determined by Two-tailed Mann-Whitney test (D), Student’s *t*-test (E), or Log-rank test (F). NS, not significant, \*\**P <* 0.01, \*\*\**P <* 0.005 and \*\*\*\**P <* 0.001 (Student’s *t*-test).

Subsequently we investigated whether the interaction of CrTir with CBL-C affects *C*. *rodentium* pathogenesis by creating *C. rodentium* Δ*tir*, and its complementary strains Δ*tir*/*tir* WT and Δ*tir*/*tir* Y451F. The series of *C*. *rodentium* strains were orally administered to C3H/HeJ mice which were then examined for bacterial load, body weight and survival rate. We sacrificed mice at 7-days post infection and measured bacterial load in the colon. The results showed that the colony forming units (CFUs) of *C. rodentium* Δ*tir*/*tir* Y451F were significantly higher than that of WT, Δ*tir*, or Δ*tir*/*tir* WT (Figure 6D). The body weight of mice infected with *C. rodentium* Δ*tir*/*tir* Y451F was drastically decreased compared to Δ*tir*/*tir* WT (Figure 6E). Finally, = mice infected with *C. rodentium* Δ*tir*/*tir* Y451F all (n=6) died by 16-days post infection, but mice infected with Δ*tir*/*tir* WT survived up to 21-days post infection (Figure 6F). These data corroborated our notion that the capacity of CBL-C to degrade Tir is substantial for attenuation of EPEC pathogenesis.

## Discussion

One of the hallmarks of EPEC infection is induction of effacement of the brush border microvilli, intimate adherence of bacteria to the epithelial cell surface, and formation of actin pedestal structure (Frankel and Phillips, 2008; Mundy et al., 2005). During EPEC infection of epithelial cells, translocated Tir acts as a versatile protein by interacting with a number of host proteins through its N- and C-terminal cytoplasmic domains, leading to actin nucleation and formation of pedestal structure beneath bacterial adherence site (Lai et al., 2013). Therefore, in the present study we sought to study the fate of Tir protein and its correlation with bacterial colonization during EPEC infection.

We report for the first time that EPEC Tir protein undergoes UPS-dependent degradation by the host CBL-C E3 ubiquitin ligase during bacterial colonization of the epithelial cells, resulting in decreased levels of cytosolic Tir. Consequently, the degradation of Tir also affects the actin pedestal structure together with the level of bacterial colonization of epithelial cells. *In vitro* analyses with GST-pull downs and immunoprecipitation showed the capability of CBL-C to directly interact with Tir in a tyrosine phosphorylation-dependent manner. Consistent with these results, *in vitro* analysis using HCT116 cells or HEK293T cells indicated that intracellular levels of Tir protein during EPEC infection greatly decreased by 3 hours to less than 10% of the original level. The decrease of Tir in HCT116 or HEK293T cell was rescued by treating cells with MG132, suggesting Tir undergoes UPS-mediated degradation (Figures 1 and S2). We also showed the tyrosine CBL-C Gly276 in the TKB domain interacts with tyrosine phosphorylated Tir (Figure 2). These results together led us to speculate that CBL-C E3 ubiquitin ligase activity acts as an important host determinant for the fate of intracellular Tir protein during EPEC infection of epithelia cells.

CBL-C is a member of the CBL E3 ubiquitin ligase family, and can negatively regulate receptor tyrosine kinase (RTK) signaling by ubiquitinating and down- regulating activated RTKs (Mohapatra et al., 2013). Compared to other CBL kinase family members such as CBL and CBL-B, CBL-C lacks C-terminus large portion containing a ubiquitin-associated (UBA) domain of CBL and CBL-B, and differs in length and amino acid sequence (Figure 2C). Previous studies using *Cbl* and *Cblb* knockout mice demonstrated that *Cbl* and *Cblb* play key physiological roles in tumor suppressors and autoimmune disease (Murphy et al., 1998; Naramura et al., 1998). However, a later study reported that *Cblc*-deficient mice were viable, healthy, and fertile and displayed no histological abnormalities up to 1.5 years of age (Griffiths et al., 2003). It was also indicated that *CBLC* tissue expression is predominant in gastrointestinal tracts, and the loss of *Cblc* was not affected by proliferation of gastrointestinal epithelium. Hence the exact role of CBL-C in the gut remains unclear at present. Since epithelium of the cecum and colon are the place where Tir-expressing enterobacteria such as EPEC prefer to attach and colonize, it is likely that the host equips itself with CBL-C to protect against attacks by intruders of pathogenic bacteria such as EPEC at localized sites of infection. In this study, we demonstrate for the first time that CBL-C has an important biological function in degradation of EPEC Tir protein. Our findings shed light on the biological function of CBL-C as a critical host defense mechanism against enteric pathogen like EPEC in the intestinal tract.

It may be worth mentioning that Tir phosphorylated on Tyr 474 binds the cellular adaptor NCK, triggering actin polymerization (Kalman et al., 1999; Kenny, 1999), though the molecular mechanisms involved in Tir phosphorylation are still under debate. Swimm et al. reported that EPEC infection of cells knocked down with *Src*, *Yes*, and *Fyn* kinases can also form pedestals (Swimm et al., 2004). They showed that in addition to SRC, YES, and FYN; some redundant kinases including ABL and ARG were incorporated in pedestal structure formation. These results seem to be different from what was published by Phillips et al, that SRC family kinase FYN was essential for Tir phosphorylation and NCK-dependent pedestal formation (Phillips et al., 2004). They used a dual bacterial system: HA-tagged Tir-expressing EPEC lacking Tir and Intimin, to transfer Tir to the cell, and Intimin-expressing *E*. *coli* to cluster Tir. They reported weak Tir Y474F phosphorylation observed *in vitro* with FYN, suggesting that FYN may be involved in phosphorylation of other tyrosine residues of Tir. As shown in Figures 4B and 4C, our *in vitro* kinase assay revealed EPEC Tir Y474F, but not Y454F, was strongly phosphorylated in the presence of FYN. Pathogenic bacterial infection is known to induce a signal cascade in which multiple kinases are involved in a highly time-coordinated manner, during the stages of infection (Chichirau et al., 2019). Therefore, it is important to understand how SRC family kinases are involved in activation of signaling cascades and thereby the fate of EPEC colonization of the intestinal epithelium.

Regarding the biological impact of EPEC Tir degradation by CBL-C on host defenses against bacterial infection, our data with cultured epithelial cells strongly supported the notion of CBL-C acting as a host safeguard against EPEC infection. This theory was strengthened by data from mice infection experiments using *C*. *rodentium* expressing Tir Y451F (Δtir/tir Y451F, CBL-C-insensitive). As shown in Figure 6D, *C*. *rodentium* expressing Tir Y451F infection of C3H/HeJ mice had higher bacterial burden in the colon at 7-days post infection than did mice infected with *C*. *rodentium* expressing WT Tir (Δtir/tir WT, CBL-C-sensitive). When mice were infected with *C*. *rodentium* expressing Tir Y451F, they gradually lost body weight and died by 16 days post-infection, while mice infected with *C. rodentium* possessing WT-Tir survived until 21 days post-infection (Figure 6F). These results corroborated that EPEC expressing CBL-C-insensitive Tir (Y451F) become more pathogenic than strains expressing CBL- C-sensitive Tir.

As EPEC induced effacement of microvilli on epithelial cells and Tir-mediated F-actin pedestal formation on polarized epithelia are aberrant physiological cellular events, we believe that elimination of Tir protein by CBL-C together with the bacterial replicative scaffold during EPEC infection must be beneficial for host cells. If our premise would be true, it may be worth developing a novel class of drugs for A/E pathogenic *E*. *coli*, which specifically target the CBL-C-Tir axis.

## Acknowledgments

The authors are grateful to Kim Minsoo for technical advice and useful discussions. We also thank Arpana Sood for English proofreading, Asaomi Kuwae and Eisuke Kuroda for technical support, and the members of the Mimuro laboratory for valuable discussions. The study was supported in part by Grant-in-Aid for Scientific Research from the Ministry of Education, Culture, Sports, Science, and Technology of Japan [17K19551, 18K07127, 19K22704 (to H.M.)], the Naito Foundation, and the Smoking Research Foundation (to H.M.).

## Author Contributions

Conceptualization, J.R., C.S. and H.M.; Methodology. J.R., H.M. H.A., and A.A.; Investigation, J.R., R.O., T.I., and H.M.; Writing–Original Draft, J.R. and H.M.; Writing– Review & Editing, C.S., and H.M.; Funding Acquisition, H.M.

## STAR Methods

### Bacterial strains

EPEC 2348/69 was used as the wild type (WT) (Nagai et al., 2005). EPEC Δ*tir* strain was kindly provided by Dr. Abe (Kitasato University). *Citrobacter rodentium* DBS100 used as the wild type (Nagai et al., 2005). The *C. rodentium* Δ*tir* strain was generated by the Red Disruption System (Datsenko and Wanner, 2000).

### Antibodies and materials

Anti-GST-Tag (91G1), anti-Myc (9B11), anti-HA (6E2), anti-lysine48-linkage specific polyubiquitin, anti-lysine63-linkage specific polyubiquitin, anti-ubiquitin P4D1 and anti-actin antibodies were purchased from Cell Signalling Technology (Danvers, MA, USA). Anti-FYN was purchased from Santa Cruz Biotechnology (Dallas, TX, USA). Anti-FLAG M2 (F3156), anti-FLAG (F7425) and anti-phosphotyrosine (4G10) were purchased from Sigma Aldrich (St. Louis, MO, USA). Anti-EspB, EspF, and Map rabbit polyclonal antibodies were used as described previously (Iizumi et al., 2007; Nagai et al., 2005). Horseradish peroxidase (HRP)-conjugated anti-mouse IgG and HRP-conjugated anti-rabbit IgG were purchased from Sigma-Aldrich. Alkaline phosphatase (AP)-conjugated anti-mouse IgG and AP-conjugated anti-rabbit IgG were purchased from Santa Cruz Biotechnology. Adenosine 5’-triphosphate (ATP) was purchased from Oriental Yeast Co. (Tokyo, Japan). Cycloheximide was purchased from Sigma-Aldrich. MG132 and Lactacystin (Peptide Inst) were obtained commercially.

### DNA constructs

The full-length human *CBL* family cDNA were cloned and introduced into pcDNA-HA. These cDNAs were then cloned into pME18S-FLAG. Mutation of Cys351 to Ala, C351A and Gly276 to Glu, *CBLC* G276E nucreotide substitutions were introduced into the human *CBLC* cDNA by site-directed mutagenesis. The human *CBLC* sequence encoding amino-acid residues 1–350, designated ΔTKB, was generated by *Bgl*II- mediated deletion. These cDNAs were then cloned into pME18S-FLAG, pME18-GST plasmids. The mouse *Cblc* coding sequence was amplified by PCR and subcloned into pME18S-FLAG. Mutation of Cys351 to Ala, C351A were introduced into the mouse *Cblc* cDNA by site-directed mutagenesis. EPEC Tir coding sequence was amplified by PCR and subcloned into pET28a, pcDNA3-Flag, pcDNA3-HA, pcDNA3-6×Myc or pME18-GST. Tyrosine residue 454, 474 in Tir was replaced with Phenylalanine by site-directed mutagenesis to generate Tir Y454F, Y474F and Y454F/Y474F in pET28a. pcDNA3-His-Ubiquitin was described previously (Otsubo et al., 2016). EPEC effector Tir coding sequence was amplified by PCR and cloned into pBluescript SK (+)-Myc and subcloned into pBR322 vector. Tir tyrosine residue 454 in Tir was replaced with Phenylalanine by site-directed mutagenesis to generate Tir Y454F. The resultant plasmid was introduced into the EPEC ΔTir strain. *C. rodentium* Tir coding sequence was amplified by PCR and cloned into pBluescript SK (+)-Myc and subcloned into pBR322 vector. Tir tyrosine residue 451 in Tir was replaced with Phenylalanine by site-directed mutagenesis to generate Tir Y451F. The resultant plasmid was introduced into the *C. rodentium* Δ*tir* strain generated by the Red Disruption System (Datsenko and Wanner, 2000). pSU-GFP vector was introduced into the EPEC WT, Δ*tir*, Δ*tir*/*tir* WT and Δ*tir*/*tir* Y454F to make GFP-EPECs.

### Cell culture

HCT116 human colon epithelial cells and HCT116 cells stably expressing pmCherry-actin were selected with 1 mg/ml G418 (Roche) was cultured in McCoy’s 5A Medium (Gibco) containing 10% fetal bovine serum. HEK293T human embryonic kidney epithelial cells were cultured in Dulbecco’s modified Eagle medium (Sigma).

### Bacterial infection

EPEC and *C. rodentium* strains were pre-cultured overnight in Luria-Bertani broth (Difco, BD, Franklin Lakes, NJ, USA) at 37°C. Bacterial cultures were inoculated into Dulbecco’s modified Eagle medium (Sigma) and incubated for 3 hours at 37°C for secretion assay. HCT116 cells were infected with the different strains of EPEC at a multiplicity of infection (MOI) of 150. After incubation for 3 hours at 37°C, plates were washed three times with PBS, transferred into fresh medium containing 100 μg/ml gentamicin and 60 μg/ml kanamycin to kill extracellular bacteria (without killing for CFU).

### Confocal microscopy analysis

HCT116 cells stably expressing pmCherry-actin were infected with GFP-EPEC. After fixation, the samples were stained with DAPI to visualize DNA. The stained samples were examined with a confocal laser-scanning microscope equipped with LSM510 version 3.2 software (LSM700; Carl Zeiss, Oberkochen, Germany).

### *In vivo* ubiquitination assay

HEK293T cells were transiently co-transfected with His-Ub, FLAG-CBL family, and HA-Tir WT or Tir Y454F. Lysates were immunoprecipitated with Ni-NTA beads and subjected to western blotting. For the proteasomal inhibitor experiments, cells were incubated with 20 μM MG132 (Peptide Inst.) at 37°C for 4hours before harvest. HCT116 cells (siRNA targeting CBL family) were transiently co-transfected with His-Ub, and HA-Tir WT. Lysates were immunoprecipitated with Ni-NTA beads and subjected to western blotting.

### Bacterial effector degradation assay

HCT116 cells with or without siRNA treatment were infected with EPEC, and were treated with 20 μM MG132, 20 μM lactacystin at 37°C for indicated time and harvested for immunoblotting. The intensity of each band was quantitated by densitometry, analysed with ImageJ software, normalized to the protein level of β-actin, and calculated in arbitrary units set to a value of 100 for uninfected cells. The results of immunoblotting experiments represent the typical datasets from three independent experiments.

### Cycloheximide chase assay

At 48 hours after transfection, HCT116 cells were treated with cycloheximide (20 μg/ml) (Wako), 20 μM MG132 or 20 μM lactacystin, and cell lysates were harvested at the indicated times for immunoblotting. The intensity of each band was quantitated by densitometry, analysed with ImageJ software, normalized with the protein level of β-actin, and calculated in arbitrary units set to a value of 100 for uninfected cells. The results of immunoblotting experiments represent the typical datasets of three independent experiments.

### RNAi

The human *CBL* family-specific siRNA sequences were as follows: CBLC1 5’-GCUCUGGGGACUUUCUACUCA-3’ CBLC2 5’- AGAGCUGCACGCACUCUUCC-3, CBLC3 5’- GAGUUCGACGUCUUCACCAGG-3 CBLC4 5’-CAGGGCACCCAGAUGUGCUGC-3 CBLC5 5’-GUUGUGAAACCGAAAUAAACU-3 CBLB1 5’- GAUCGUGAGGAGUCCUUGACG-3 CBLB2 5’-GGAGACCGAUGCUUGCUCAGG-3 CBLB3 5’-CAGGGUUGAACUGACAAAAAU-3 CBL1 5’- CAGCAUCUCCGUACUAUCUUG-3 CBL2 5’-CGAGUUCAGUUAGUAUUCUGU-3 CBL3 5’-GCGGUCCCGAGUUGAGGUAGA-3 SiLuc 5’CGUACGCGGAAUACUUCGAdTdT (Sigma). Cells were transfected using RNAiMax (Invitrogen). After 72 hours, siRNA-treated cells were used for further analysis

### Real-time RT-PCR analysis

Total RNAs was prepared with ISOGEN (Nippon gene), and cDNAs was obtained by reverse transcription using Superscript II reverse transcriptase (Invitrogen) and amplified by PCR. Primer pairs were as follows; *CBLC* 5’- AGGATGTGAAGATTGAGCCG-3’ 3’-GCCGTGAGTATCTACCAGTTC-5’, *CBLB* 5’- CAGGAACAATATGAATTATA-3’, 3’-TGCCGATGCTAGACTTGGAC-5’, *CBL* 5’- CAGGAACAATATGAATTATA-3’, 3’-TGCCGATGTGAAATTAAAGG-5 (Invitrogen).

Expression levels were normalized to the levels of HPRT.

### Purification of proteins

The genes encoding Tir were amplified by PCR into the pET28a vector. Site-directed point mutagenesis of Tir was done by PCR. Sequence encoding histidine fusion proteins was expressed in BL-21(DE3) *E. coli* and purified by Ni-NTA agarose beads (QIAGEN). Afterward, proteins were dialyzed against a buffer of 20 mM Tris-HCl (pH 7.5), 5 mM MgCl2 and 1 mM DTT. For production of bacuroviral GST-FYN protein, cDNAs for *FYN* was cloned into the pFastBAC-GST and Purified. GST was removed from GST–FYN with PreScission protease (GE Healthcare) treatment.

### GST pull-down assay

HEK293T cells were transiently transfected with plasmids using FuGENE 6 (Roche). Cells were washed with PBS and lysed for 30 min at 4°C in lysis buffer containing 150 mM NaCl, 50 mM Tris-HCl (pH 7.5), 1 mM EDTA, 0.5% Triton X-100, and Complete Protease Inhibitor Cocktail (Roche). Lysates were cleared by centrifugation, and Glutathione S-transferase (GST)-fused proteins were pull-downed for 2 hours with Glutathione Sepharose 4B beads (GE) at 4°C. The bound proteins were washed five times with lysis buffer and subjected to western blotting.

### Immunoprecipitation

HEK293T cells were transiently transfected with plasmids using FuGENE 6 (Roche). Cells were washed with PBS and lysed for 30 min at 4°C in lysis buffer containing 150 mM NaCl, 50 mM Tris-HCl (pH 7.5), 1 mM EDTA, 0.5% Triton X-100, and Complete Protease Inhibitor Cocktail (Roche). Lysates were cleared by centrifugation, and FLAG-tagged proteins were immunoprecipitated for 2 hours with anti-FLAG M2 and protein G beads (Sigma) at 4°C. Immunoprecipitates were washed five times with lysis buffer and subjected to western blotting.

### *In vivo* ubiquitination assay

HEK293T cells were transiently co-transfected with His-Ub, FLAG-CBL family or CBL- C CA, and HA-Tir WT, Tir Y454F, *Citrobacter rodentium* HA-Tir WT or Tir Y451F. The lysates were pulled down with Ni-NTA beads. Bound proteins and cell lysates were subjected to western blotting.

### *In vitro* kinase assays

His-Tir and derivatives were expressed in BL21(DE3) and purified by Ni-NTA beads. Tyrosine Kinase proteins and FYN proteins were expressed in baculovirus-infected Sf9 insect cells and purified. GST-FYN was purified and the GST was removed. *In vitro* kinase reaction were performed as described (Nakazawa et al., 2001).

### Preparation of Proteins from Culture Supernatant

Proteins released into bacterial culture supernatants were concentrated by trichloroacetic acid (TCA) precipitation. The culture supernatants were filtered and trichloroacetic acid was then added to each sample at a final concentration of 10% (v/v). After incubation on ice for 1- hour, the samples were centrifuged for 5 min. The resulting precipitated proteins were dissolved in the SDS-PAGE sample buffer.

### Animal protocol

Oral infection, tissue collection and CFU counts. *C. rodentium* strain DBS100, its Δ*tir*/*tir* WT and Δ*tir*/*tir* Y451F mutants were grown in LB broth by shaking overnight at 37 °C. 6 to 8-week-old female C3H/HeJ or C57BL/6 mice (Japan SLC, Inc.) were orally administered with 2.5 × 10^8^ CFU of the indicated *C. rodentium* strains in a total volume of 200 µl per mouse. Mortality and body weights were monitored daily after infection.

For CFU counts, mice were sacrificed at 7 (C3H/HeJ) to 12 (C57BL/6) days after infection, colons were removed aseptically, weighed and homogenized in PBS. The homogenates were serially diluted and plated on MacConkey agar plates. Bacterial colonies were counted after overnight incubation at 37 °C. All animal experiments were conducted in accordance with the guidelines of the University of Tokyo for the care and use of laboratory animals and were approved by the ethics committee for animal experiments of the University of Tokyo.

### Statistical analysis

Statistically significant differences between mean values were determined using Prism6 (GraphPad Software Inc., La Jolla, CA, USA). Data are presented as the means ± SD or SEM. A P-value of P < 0.05 was considered to indicate a statistically significant difference.

## Supplemental Information

**Figure S1.**
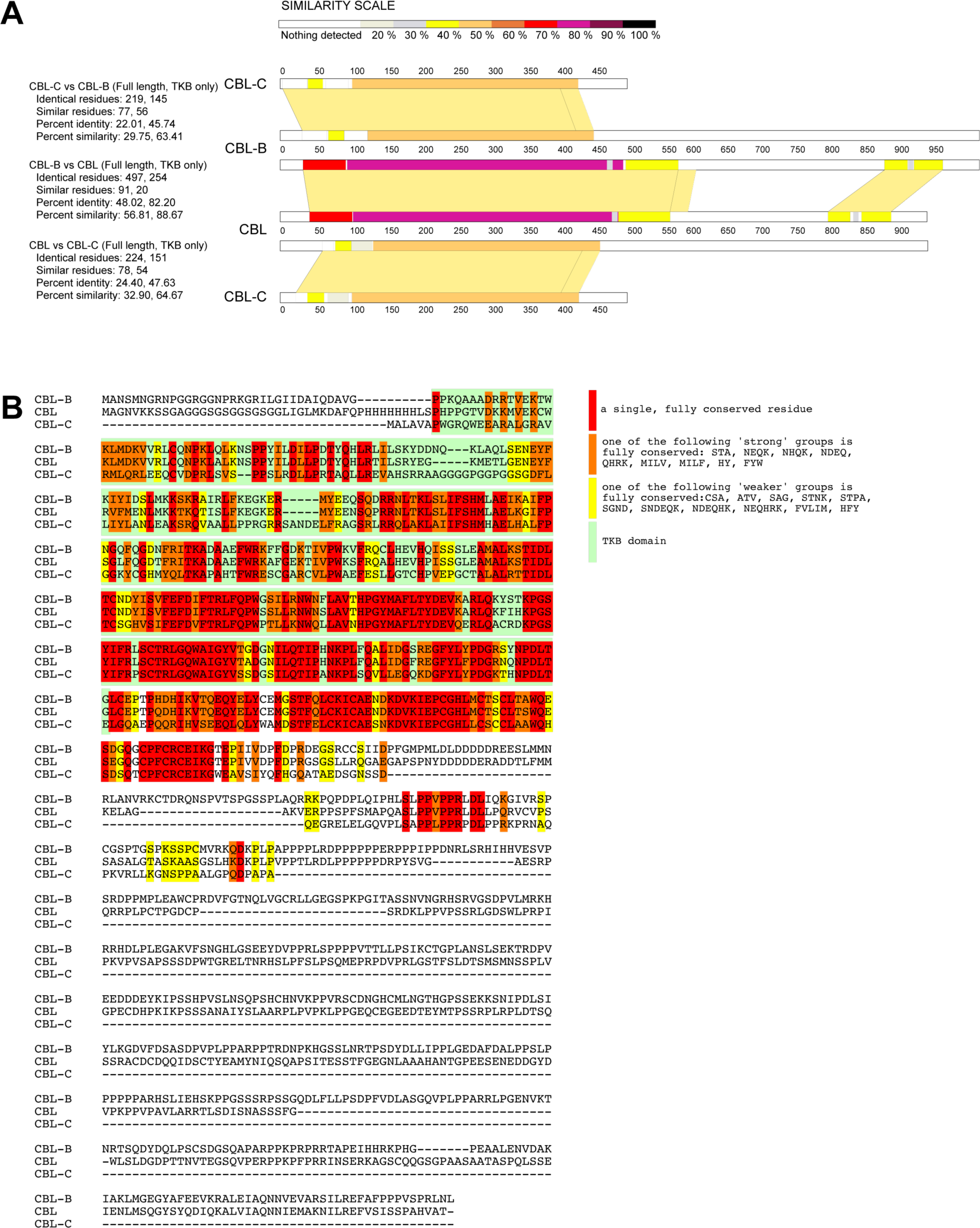
Related to Figure 2. **(A)** Amino acid sequence alignments of CBL, CBL-B, and CBL-C. Each percent identity and percent similarity were analyzed using COBALT Constraint-based Multiple Alignment Tool (https://www.ncbi.nlm.nih.gov/tools/cobalt/cobalt.cgi?CMD=Web) and ExPASy SIM analysis (https://web.expasy.org/sim/), and similarity scales were illustrated using LALNVIEW version lalnview 3.0. **(B)** Amino acid sequence alignments of CBL, CBL-B, and CBL-C. Protein-protein BLAST and ClustalW2 tools were used for the analysis.

**Figure S2.**
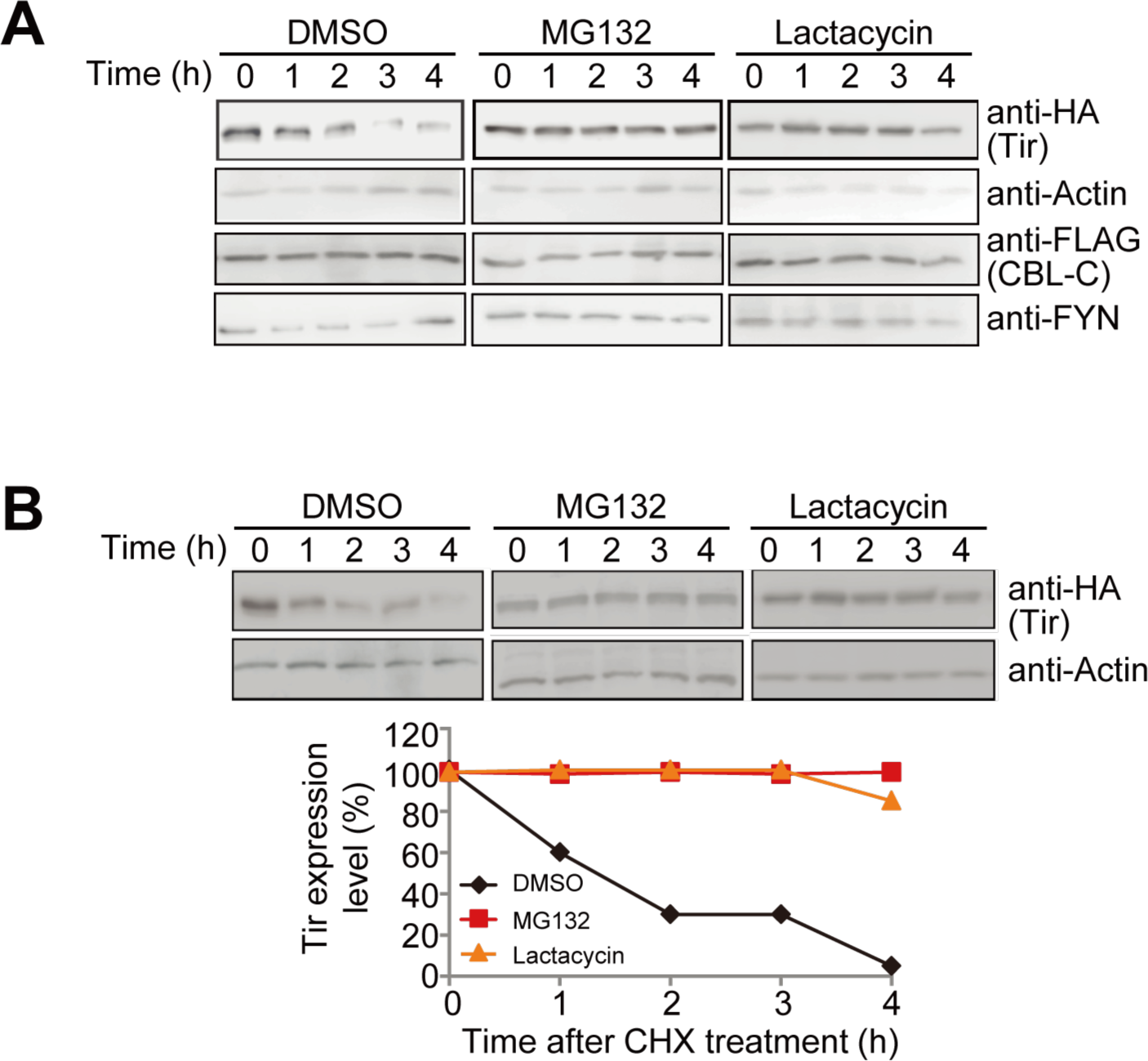
Related to Figure 3. **(A)** HEK293T cells or **(B)** HCT116 cells were transfected with indicated plasmids. At 48 hours after transfection, cells were treated with DMSO, MG132 (20 μM) or Lactacystin (20 μM) and 10 μM of cycloheximide (CHX). After incubation for the indicated time, cells were harvested and lysates were separated by SDS-PAGE and subjected to western blot with the noted antibodies. The intensity of Tir protein was normalized to the intensity of β-actin (Actin) and calculated in arbitrary units set to a value of 100 % for 0 hour. These results represent typical examples of three separate experiments. Densitometric analysis was conducted using ImageJ software.

**Figure S3.**
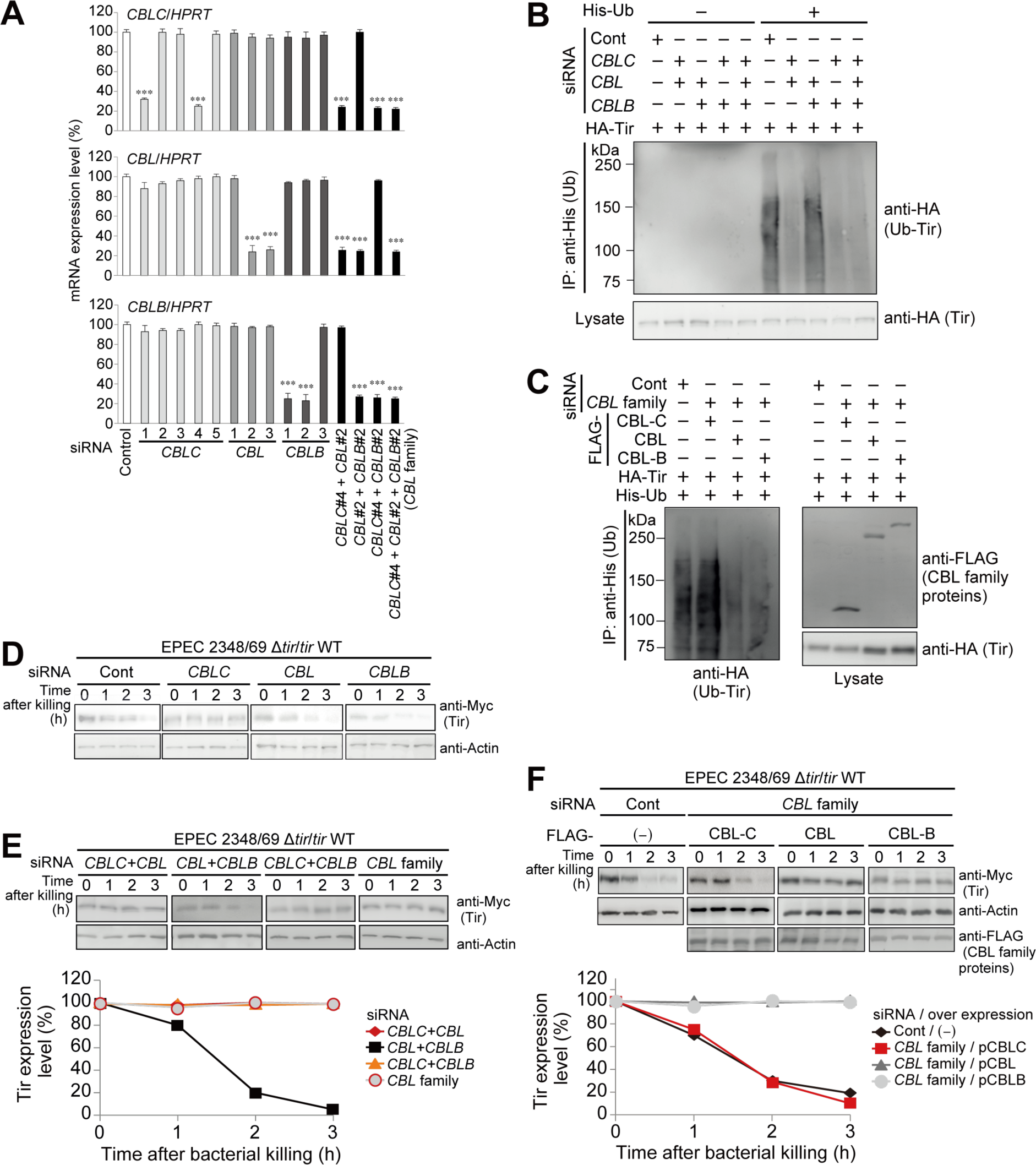
Related to Figure 3. (A) HCT116 cells were transfected with siRNA targeting *CBL* family or Luc (control). At 72 hours of transfection, the lysates were harvested and subjected to quantitative RT-PCR analysis of *CBLC*, *CBL* and *CBLB*. *CBLC* #4, *CBL* #2, and *CBLB* #2 were used in subsequent experiments. Data are representative of three independent experiments (mean and SEM). n=xx, \*\*\**P <* 0.001 vs control (Student’s *t*-test). (B) HCT116 cells were transfected with siRNA targeting CBL family or Luc (control), then transfected with the indicated plasmids. The lysates were pulled down with Ni-NTA beads. Bound proteins were separated by SDS-PAGE and subjected to western blot with the noted antibodies. (C) HCT116 cells transfected with siRNA targeting *CBL* family (including *CBLC*, *CBL*, and *CBLB*) or Luc (control) were subjected to additional transfection of the indicated plasmids. The cells were washed 3 times with PBS and lysed for 30min at RT in lysis buffer. Lysates were cleared by centrifugation, and proteins were pulled down for 2 hours with Ni-NTA beads at RT. Bound proteins were washed five times with lysis buffer, separated by SDS-PAGE and subjected to western blot with the noted antibodies. (**D and E**) HCT116 cells transfected with indicated siRNA targeting *CBL* family or Luc (control) were infected with EPEC Δ*tir*/*tir* WT at a MOI of 150 for 3 hours. The cells were washed 3 times with PBS, and then transferred into fresh medium containing gentamicin and kanamycin to kill extracellular bacteria. After incubation for the indicated time, the cells were harvested and lysates separated by SDS-PAGE and subjected to western blot with the noted antibodies. The intensity of Tir protein was normalized to the intensity of β-actin (Actin) and calculated in arbitrary units set to a value of 100 % for 0 hour after killing (E). Densitometric analysis was conducted using ImageJ software. (**F**) HCT116 cells transfected with siRNA targeting *CBL* family or Luc (control) were subjected to additional transfection of the indicated FLAG-tagged CBL family. The cells were infected with EPEC Δtir/tir WT at a MOI of 150 for 3 hours. The cells were washed 3 times with PBS, and then transferred into fresh medium containing gentamicin and kanamycin to kill extracellular bacteria. After incubation for the indicated time, cells were harvested and lysates separated by SDS-PAGE and subjected to western blot with the noted antibodies. The intensity of Tir protein was normalized to the intensity of β-actin (Actin) and calculated in arbitrary units set to a value of 100 % for 0 hour after killing. Densitometric analysis was conducted using ImageJ software.

**Figure S4.**
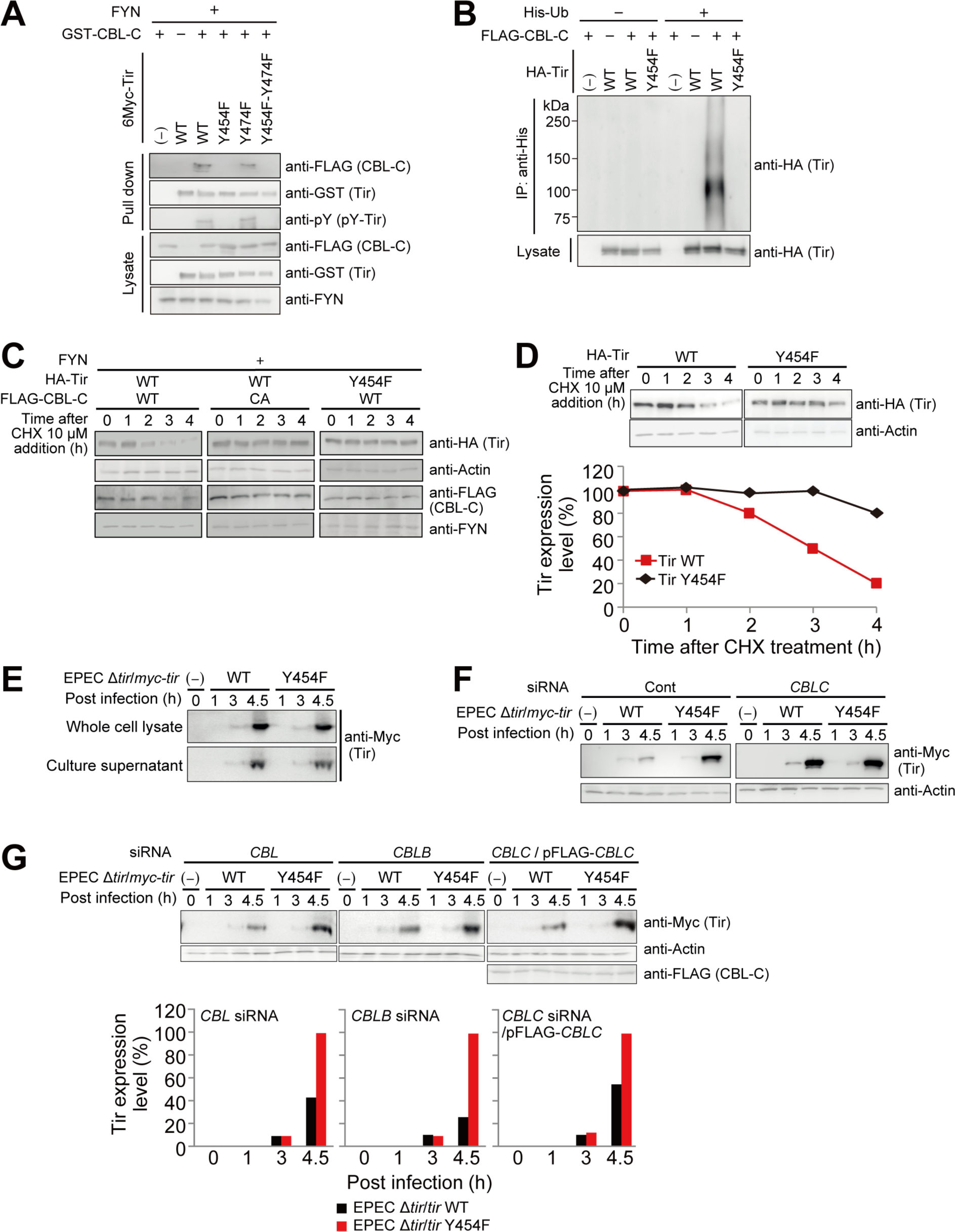
Related to Figure 4 (A) CBL-C binds to phosphorylated Tir at residue Tyrosine 454. HEK293T cells were transiently transfected with indicated plasmids. The cells were washed 3 times with PBS and lysed for 30 minutes at 4°C in lysis buffer. Lysates were cleared by centrifugation and proteins were pulled down for 2 hours with Glutathione Sepharose 4B beads at 4°C. Bound proteins were washed 5 times with lysis buffer, separated by SDS-PAGE, and subjected to Western blot with the described antibodies. (B) Tir residue Tyrosine 454 is important for Tir ubiquitination. HCT116 cells were transiently transfected with the indicated plasmids combinations. Cells were washed 3 times with PBS and lysed in lysis buffer for 30 minutes at RT. Lysates were cleared by centrifugation, and proteins were pulled down with Ni-NTA beads for 2 hours at RT. Bound proteins were washed 5 times with lysis buffer, separated by SDS-PAGE, and subjected to Western blot with the indicated antibodies. These results represent the typical examples of 3 separate experiments. (**C, D**) Tir residue Tyrosine 454 is important for Tir degradation HEK239T (C) or HCt116 (D) cells expressing the indicated plasmids were treated with chcloheximide (CHX, 10 μM). After incubation for the indicated time, cells were harvested and lysates separated by SDS-PAGE and subjected to Western blot with the indicated antibodies. The intensity of Tir protein was normalized to the intensity of β-actin (Actin) and calculated in arbitrary units set to a value of 100 % for 0 hour, The results represent the typical examples of 3 separate experiments. Densitometric analysis was conducted using ImageJ software. (**E**) Y454F Tir protein levels in EPEC Δ*tir*/*tir* Y454F are equal to that in EPEC Δ*tir*/*tir* WT. Bacterial whole cell lysates (Bacteria) and culture supernatants (Medium) were separated by SDS-PAGE and subjected to western blot with the noted antibodies. The intensity of Tir protein was calculated in arbitrary units set to a value of 100 % for samples of EPEC Δ*tir*/*tir* WT at 4.5 hours. These results represent typical examples of three separate experiments. Densitometric analysis was conducted using ImageJ software. (**F** and **G**) CBL-C regulate protein levels of intracellular Tir secreted by EPEC. HCT116 cells transfected with indicated siRNA targeting *CBL* family or Luc (control) were subjected to additional transfection of CBL-C protein, and were infected with the indicated EPEC strains at a MOI of 150 for indicated time. The cell lysates were harvested and the lysates were separated by SDS-PAGE and subjected to western blot with the noted antibodies. The intensity of Tir protein was normalized to the intensity of β-actin (Actin). The intensity of Tir protein was calculated in arbitrary units set to each value of 100 % for samples of EPEC Δ*tir*/*tir* Y454F at 4.5 hours. These results represent typical examples of three separate experiments (mean and SEM). Densitometric analysis was conducted using ImageJ software.

**Figure S5.**
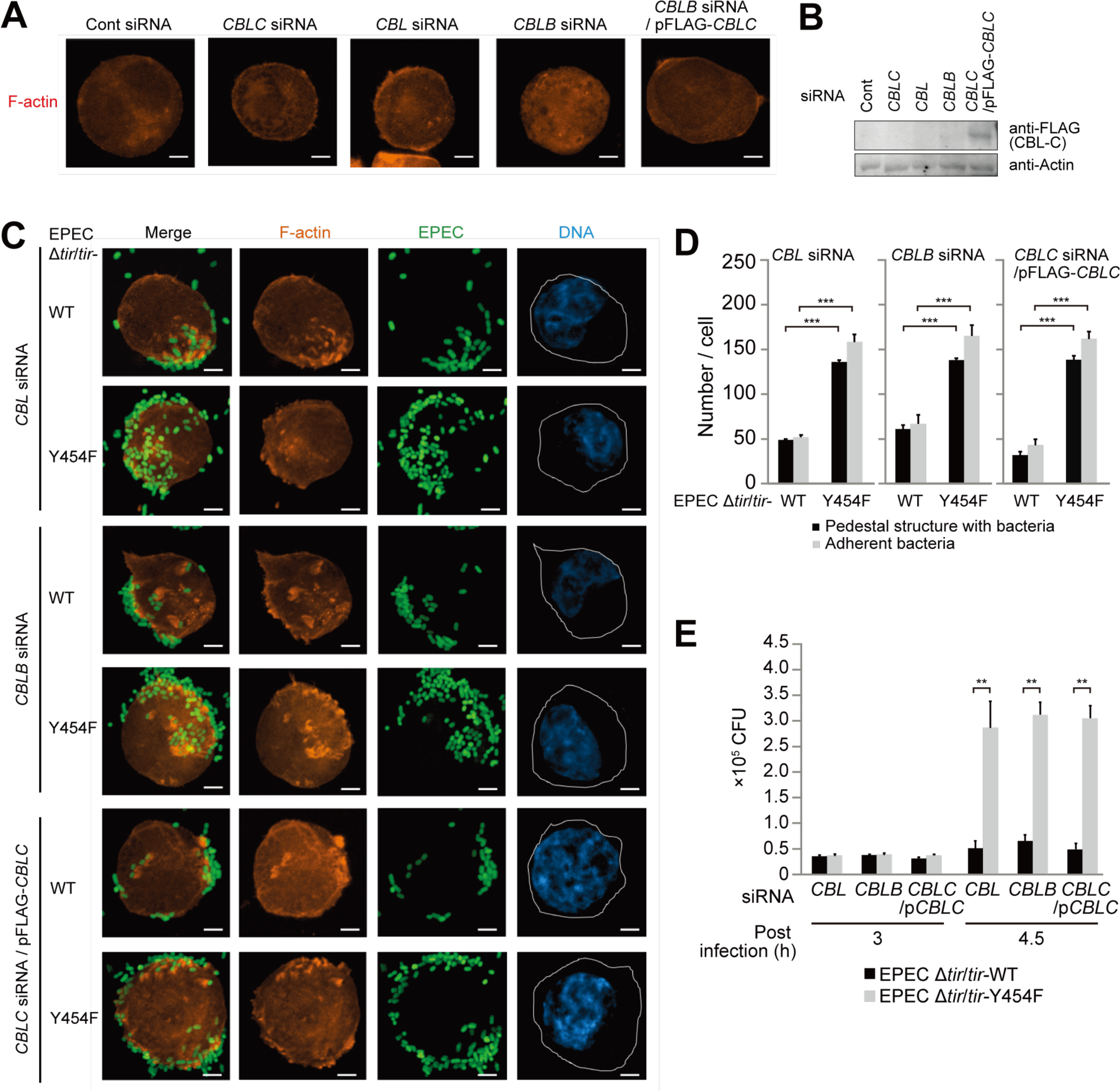
Related to Figure 5 (**A** and **B**) pmCherry actin stably expressing-HCT116 cells were transfected with siRNA targeting *CBL* family or Luc (control), and transfected with CBL-C expression plasmid (pCBL-C). Cells were fixed and actin was visualized (orange). Scale bar, 5 µm (A). (**C** and **D**) HCT116 cells transfected with siRNA targeting *CBL* family or Luc (control), or subjected to additional transfection of CBL-C protein were infected with the indicated GFP-EPEC strains at a MOI of 150 for 4.5 hours. The cells were fixed and actin (red), EPEC (green), and DNA (stained with DAPI, blue) were visualized. Scale bar, 5 µm (C). Quantification of pedestal structure and adhesion bacteria are shown (D). These results represent typical examples of three separate experiments. Data are represented as mean ± SEM (n = 150). \**P <* 0.01, \*\**P <* 0.005 and \*\*\**P <* 0.001 (Student’s *t*-test). (**E**) HCT116 cells were transfected with siRNA of each *CBLC* family or Luc (control), and infected with the indicated EPEC strains at a MOI of 150 for indicated time. The adhered bacteria were harvested at the indicated time points and plated on LB agar-plates to count the CFU. These results represent typical examples of three separate experiments. Data are represented as mean ± SEM (n = 3). \**P <* 0.01, \*\**P <* 0.005 and \*\*\**P <* 0.001 (Student’s *t*-test).

## References

Brady, M.J., Campellone, K.G., Ghildiyal, M., and Leong, J.M. (2007). Enterohaemorrhagic and enteropathogenic Escherichia coli Tir proteins trigger a common Nck-independent actin assembly pathway. Cell Microbiol 9, 2242–2253.

Campellone, K.G., and Leong, J.M. (2005). Nck-independent actin assembly is mediated by two phosphorylated tyrosines within enteropathogenic Escherichia coli Tir. Mol Microbiol 56, 416–432.

Campellone, K.G., Giese, N., Tipper, O.J., and Leong, J.M. (2002). A tyrosine-phosphorylated 12-amino-acid sequence of enteropathogenic Escherichia coli Tir binds the host adaptor protein Nck and is required for Nck localization to actin pedestals. Mol Microbiol 43, 1227–1241.

Chiang, Y.J., Kole, H.K., Brown, K., Naramura, M., Fukuhara, S., Hu, R.-J., Jang, I.K., Gutkind, J.S., Shevach, E., and Gu, H. (2000). Cbl-b regulates the CD28 dependence of T-cell activation. Nature 403, 216–220.

Chichirau, B.E., Diechler, S., Posselt, G., and Wessler, S. (2019). Tyrosine Kinases in Helicobacter pylori Infections and Gastric Cancer. Toxins 11, 591.

Datsenko, K.A., and Wanner, B.L. (2000). One-step inactivation of chromosomal genes in Escherichia coli K-12 using PCR products. Proc National Acad Sci 97, 6640– 6645.

Deibel, C., Krämer, S., Chakraborty, T., and Ebel, F. (1998). EspE, a novel secreted protein of attaching and effacing bacteria, is directly translocated into infected host cells, where it appears as a tyrosine-phosphorylated 90 kDa protein. Mol Microbiol 28, 463–474.

Donnenberg, M.S., Tacket, C.O., James, S.P., Losonsky, G., Nataro, J.P., Wasserman, S.S., Kaper, J.B., and Levine, M.M. (1993). Role of the eaeA gene in experimental enteropathogenic Escherichia coli infection. J Clin Invest 92, 1412–1417.

Frankel, G., and Phillips, A.D. (2008). Attaching effacing Escherichia coli and paradigms of Tir-triggered actin polymerization: getting off the pedestal. Cell Microbiol 10, 549–556.

Grice, G.L., and Nathan, J.A. (2016). The recognition of ubiquitinated proteins by the proteasome. Cell Mol Life Sci 73, 3497–3506.

Griffiths, E.K., Sanchez, O., Mill, P., Krawczyk, C., Hojilla, C.V., Rubin, E., Nau, M.M., Khokha, R., Lipkowitz, S., Hui, C., et al. (2003). Cbl-3-Deficient Mice Exhibit Normal Epithelial Development. Mol Cell Biol 23, 7708–7718.

Hicks, S., Frankel, G., Kaper, J.B., Dougan, G., and Phillips, A.D. (1998). Role of Intimin and Bundle-Forming Pili in Enteropathogenic Escherichia coli Adhesion to Pediatric Intestinal Tissue In Vitro. Infect Immun 66, 1570–1578.

Iizumi, Y., Sagara, H., Kabe, Y., Azuma, M., Kume, K., Ogawa, M., Nagai, T., Gillespie, P.G., Sasakawa, C., and Handa, H. (2007). The enteropathogenic E. coli effector EspB facilitates microvillus effacing and antiphagocytosis by inhibiting myosin function. Cell Host Microbe 2, 383–392.

Kalman, D., Weiner, O.D., Goosney, D.L., Sedat, J.W., Finlay, B.B., Abo, A., and Bishop, J.M. (1999). Enteropathogenic E. coli acts through WASP and Arp2/3 complex to form actin pedestals. Nat Cell Biol 1, 389–391.

Kenny, B. (1999). Phosphorylation of tyrosine 474 of the enteropathogenic Escherichia coli (EPEC) Tir receptor molecule is essential for actin nucleating activity and is preceded by additional host modifications. Mol Microbiol 31, 1229–1241.

Kenny, B., and Warawa, J. (2001). Enteropathogenic Escherichia coli (EPEC) Tir Receptor Molecule Does Not Undergo Full Modification When Introduced into Host Cells by EPEC-Independent Mechanisms. Infect Immun 69, 1444–1453.

Kim, M., Tezuka, T., Tanaka, K., and Yamamoto, T. (2004). Cbl-c suppresses v-Src-induced transformation through ubiquitin-dependent protein degradation. Oncogene 23, 1645–1655.

Lai, Y., Rosenshine, I., Leong, J.M., and Frankel, G. (2013). Intimate host attachment: enteropathogenic and enterohaemorrhagic Escherichia coli. Cell Microbiol 15, 1796– 1808.

Lilienbaum, A. (2013). Relationship between the proteasomal system and autophagy. International Journal of Biochemistry and Molecular Biology 4, 1 26.

Lill, N.L., Douillard, P., Awwad, R.A., Ota, S., Lupher, M.L., Miyake, S., Meissner-Lula, N., Hsu, V.W., and Band, H. (2000). The Evolutionarily Conserved N-terminal Region of Cbl Is Sufficient to Enhance Down-regulation of the Epidermal Growth Factor Receptor. J Biol Chem 275, 367–377.

Liu, H., Magoun, L., Luperchio, S., Schauer, D.B., and Leong, J.M. (1999). The Tir-binding region of enterohaemorrhagic Escherichia coli intimin is sufficient to trigger actin condensation after bacterial-induced host cell signalling. Mol Microbiol 34, 67– 81.

Meng, W., Sawasdikosol, S., Burakoff, S.J., and Eck, M.J. (1999). Structure of the amino-terminal domain of Cbl complexed to its binding site on ZAP-70 kinase. Nature 398, 84–90.

Mohapatra, B., Ahmad, G., Nadeau, S., Zutshi, N., An, W., Scheffe, S., Dong, L., Feng, D., Goetz, B., Arya, P., et al. (2013). Protein tyrosine kinase regulation by ubiquitination: Critical roles of Cbl-family ubiquitin ligases. Biochimica Et Biophysica Acta Bba - Mol Cell Res 1833, 122–139.

Mundy, R., MacDonald, T.T., Dougan, G., Frankel, G., and Wiles, S. (2005). Citrobacter rodentium of mice and man. Cell Microbiol 7, 1697–1706.

Murphy, M.A., Schnall, R.G., Venter, D.J., Barnett, L., Bertoncello, I., Thien, C.B.F., Langdon, W.Y., and Bowtell, D.D.L. (1998). Tissue Hyperplasia and Enhanced T-Cell Signalling via ZAP-70 in c-Cbl-Deficient Mice. Mol Cell Biol 18, 4872–4882.

Nagai, T., Abe, A., and Sasakawa, C. (2005). Targeting of Enteropathogenic Escherichia coli EspF to Host Mitochondria Is Essential for Bacterial Pathogenesis CRITICAL ROLE OF THE 16TH LEUCINE RESIDUE IN EspF. J Biol Chem 280, 2998–3011.

Nakazawa, T., Komai, S., Tezuka, T., Hisatsune, C., Umemori, H., Semba, K., Mishina, M., Manabe, T., and Yamamoto, T. (2001). Characterization of Fyn-mediated Tyrosine Phosphorylation Sites on GluRε2 (NR2B) Subunit of theN-Methyl-d-aspartate Receptor. J Biol Chem 276, 693–699.

Naramura, M., Kole, H.K., Hu, R.-J., and Gu, H. (1998). Altered thymic positive selection and intracellular signals in Cbl-deficient mice. Proc National Acad Sci 95, 15547–15552.

Ota, S., Hazeki, K., Rao, N., Lupher, M.L., Andoniou, C.E., Druker, B., and Band, H. (2000). The RING Finger Domain of Cbl Is Essential for Negative Regulation of the Syk Tyrosine Kinase. J Biol Chem 275, 414–422.

Otsubo, R., Kim, M., Lee, J., and Sasakawa, C. (2016). Midori-ishi Cyan/monomeric Kusabira-Orange-based fluorescence resonance energy transfer assay for characterization of various E3 ligases. Genes Cells 21, 608–623.

Phillips, N., Hayward, R.D., and Koronakis, V. (2004). Phosphorylation of the enteropathogenic E. coli receptor by the Src-family kinase c-Fyn triggers actin pedestal formation. Nat Cell Biol 6, 618–625.

Sason, H., Milgrom, M., Weiss, A.M., Melamed-Book, N., Balla, T., Grinstein, S., Backert, S., Rosenshine, I., and Aroeti, B. (2009). Enteropathogenic Escherichia coli Subverts Phosphatidylinositol 4,5-Bisphosphate and Phosphatidylinositol 3,4,5- Trisphosphate upon Epithelial Cell Infection. Mol Biol Cell 20, 544–555.

Selbach, M., Paul, F.E., Brandt, S., Guye, P., Daumke, O., Backert, S., Dehio, C., and Mann, M. (2009). Host cell interactome of tyrosine-phosphorylated bacterial proteins. Cell Host Microbe 5, 397–403.

Stevens, J.M., Galyov, E.E., and Stevens, M.P. (2006). Actin-dependent movement of bacterial pathogens. Nat Rev Microbiol 4, 91–101.

Swaminathan, G., and Tsygankov, A.Y. (2006). The Cbl family proteins: Ring leaders in regulation of cell signaling. J Cell Physiol 209, 21–43.

Swimm, A., Bommarius, B., Li, Y., Cheng, D., Reeves, P., Sherman, M., Veach, D., Bornmann, W., and Kalman, D. (2004). Enteropathogenic Escherichia coli Use Redundant Tyrosine Kinases to Form Actin Pedestals. Mol Biol Cell 15, 3520–3529.

Thien, C.B.F., and Langdon, W.Y. (2001). Cbl: many adaptations to regulate protein tyrosine kinases. Nat Rev Mol Cell Bio 2, 294–307.

Vallance, B.A., Deng, W., Jacobson, K., and Finlay, B.B. (2003). Host Susceptibility to the Attaching and Effacing Bacterial Pathogen Citrobacter rodentium. Infect Immun 71, 3443–3453.

Vingadassalom, D., Campellone, K.G., Brady, M.J., Skehan, B., Battle, S.E., Robbins, D., Kapoor, A., Hecht, G., Snapper, S.B., and Leong, J.M. (2010). Enterohemorrhagic E. coli Requires N-WASP for Efficient Type III Translocation but Not for EspFU- Mediated Actin Pedestal Formation. Plos Pathog 6, e1001056.

Walczak, H., Iwai, K., and Dikic, I. (2012). Generation and physiological roles of linear ubiquitin chains. Bmc Biol 10, 23.

Wong, A.R.C., Pearson, J.S., Bright, M.D., Munera, D., Robinson, K.S., Lee, S.F., Frankel, G., and Hartland, E.L. (2011). Enteropathogenic and enterohaemorrhagic Escherichia coli: even more subversive elements. Mol Microbiol 80, 1420–1438.

Yan, D., Wang, X., Luo, L., Cao, X., and Ge, B. (2012). Inhibition of TLR signaling by a bacterial protein containing immunoreceptor tyrosine-based inhibitory motifs. Nat Immunol 13, 1063–1071.

Zheng, N., Wang, P., Jeffrey, P.D., and Pavletich, N.P. (2000). Structure of a c-Cbl– UbcH7 Complex RING Domain Function in Ubiquitin-Protein Ligases. Cell 102, 533– 539.

